# Bias toward long gene misregulation in synaptic disorders can be an artefact of amplification-based methods

**DOI:** 10.1101/240705

**Authors:** Ayush T. Raman, Amy E. Pohodich, Ying-Wooi Wan, Hari Krishna Yalamanchili, Bill Lowry, Huda Y. Zoghbi, Zhandong Liu

## Abstract

Several recent studies have suggested that genes that are longer than 100 kilobases are more likely to be misregulated in neurological diseases associated with synaptic dysfunction, such as autism and Rett syndrome. These length-dependent transcriptional changes are modest in *Mecp2*-mutant samples, but, given the low sensitivity of high-throughput transcriptome profiling technology, the statistical significance of these results needs to be re-evaluated. Here, we show that the apparent length-dependent trends previously observed in MeCP2 microarray and RNA-Sequencing datasets, particularly in genes with low fold-changes, disappeared after accounting for baseline variability estimated from randomized control samples. As we found no similar bias with NanoString technology, this long-gene bias seems to be particular to PCR amplification-based platforms. In contrast, authentic long gene effects, such as those caused by topoisomerase inhibition, can be detected even after adjustment for baseline variability. Accurate detection of length-dependent trends requires establishing a baseline from randomized control samples.

**HIGHLIGHTS:**

- Length-dependent gene misregulation is not intrinsic to *Mecp2* disruption.
- Topoisomerase inhibition produces an authentic long gene bias.
- PCR amplification-based high-throughput datasets are biased toward long genes.

## INTRODUCTION

The capacity for large-scale analysis of transcriptional changes in human disease has attracted considerable research attention, most recently in studies related to autism spectrum disorders, including Angelman syndrome, Rett syndrome (RTT), Fragile X syndrome, and autism itself (Zoghbi and Bear, 2012). Microarray and RNA-Seq studies have demonstrated that these disorders involve the dysregulation of thousands of neuronal genes. Several recent studies have also suggested that the genes dysregulated in these syndromes tend to be those that consist of more than 100 kilobases (Katz et al., 2016; Zylka et al., 2015). This intriguing length bias has been observed across both epigenetic and transcriptional datasets such as Angelman syndrome (Huang et al., 2011), Rett syndrome (Gabel et al., 2015; Kinde et al., 2016; Sugino et al., 2014), Fragile X syndrome (Gabel et al., 2015; Ouwenga and Dougherty, 2015) autism (King et al., 2013; Sullivan et al., 2015). The degree of bias tends to be fairly mild, however, long genes are themselves overrepresented in the brain compared to other tissues in the body (Zylka et al., 2015). It seems worthwhile to examine this apparent bias more closely in gene expression datasets.

The afore-mentioned gene expression studies (Gabel et al., 2015; King et al., 2013; Sugino et al., 2014; Sullivan et al., 2015) partitioned the entire genome into hundreds of overlapping bins (or windows), with each bin containing hundreds of genes. Within each bin, the average fold-change in wildtype or untreated brain tissue was compared to that observed in the knock-out or treatment groups, and a running average log2 fold-change was plotted against the average gene length. In these running average plots, long genes demonstrated a non-zero mean compared to short genes. These analyses did not, however, establish a baseline of inherent variation among samples within a given genotype, and they did not employ a statistical test to determine the significance of the length-dependent changes. It should be noted that variations in measured gene expression can arise because of RNA priming (Hansen et al., 2010; Li et al., 2010), GC-content (Risso et al., 2011), transcript length (Oshlack and Wakefield, 2009), or library preparation (Lahens et al., 2014), all of which must be accounted for in order to avoid unwarranted biological conclusions (Robert and Watson, 2015; Wan et al., 2014).

We, therefore, analysed a comprehensive list of large datasets derived from different transcriptome profiling technologies and set out to determine the best way to enhance the signal-to-noise ratio. To this end, we began by analysing technical replicates using benchmark datasets. Using these datasets, we developed an approach to reliably identify patterns with respect to gene regulation, and we then applied our approach to analyse datasets for which long gene trends have been reported.

## RESULTS

### Baseline length dependency should first be estimated from the control groups: the Topotecan study as a positive control

Preferential dysregulation of long genes is generally estimated by computing the average gene expression fold-changes between experimental groups and plotting this fold-change against the gene length (Gabel et al., 2015; King et al., 2013; Sugino et al., 2014), also known as running average plots (red curve in Fig 1A, Experimental Procedures). However, the statistical significance of running average plots has never been evaluated in the current literature. Here, we propose an approach to estimate statistical significance by constructing a null distribution of the running average plot from randomized control samples (Figure S1).

**Figure 1.**
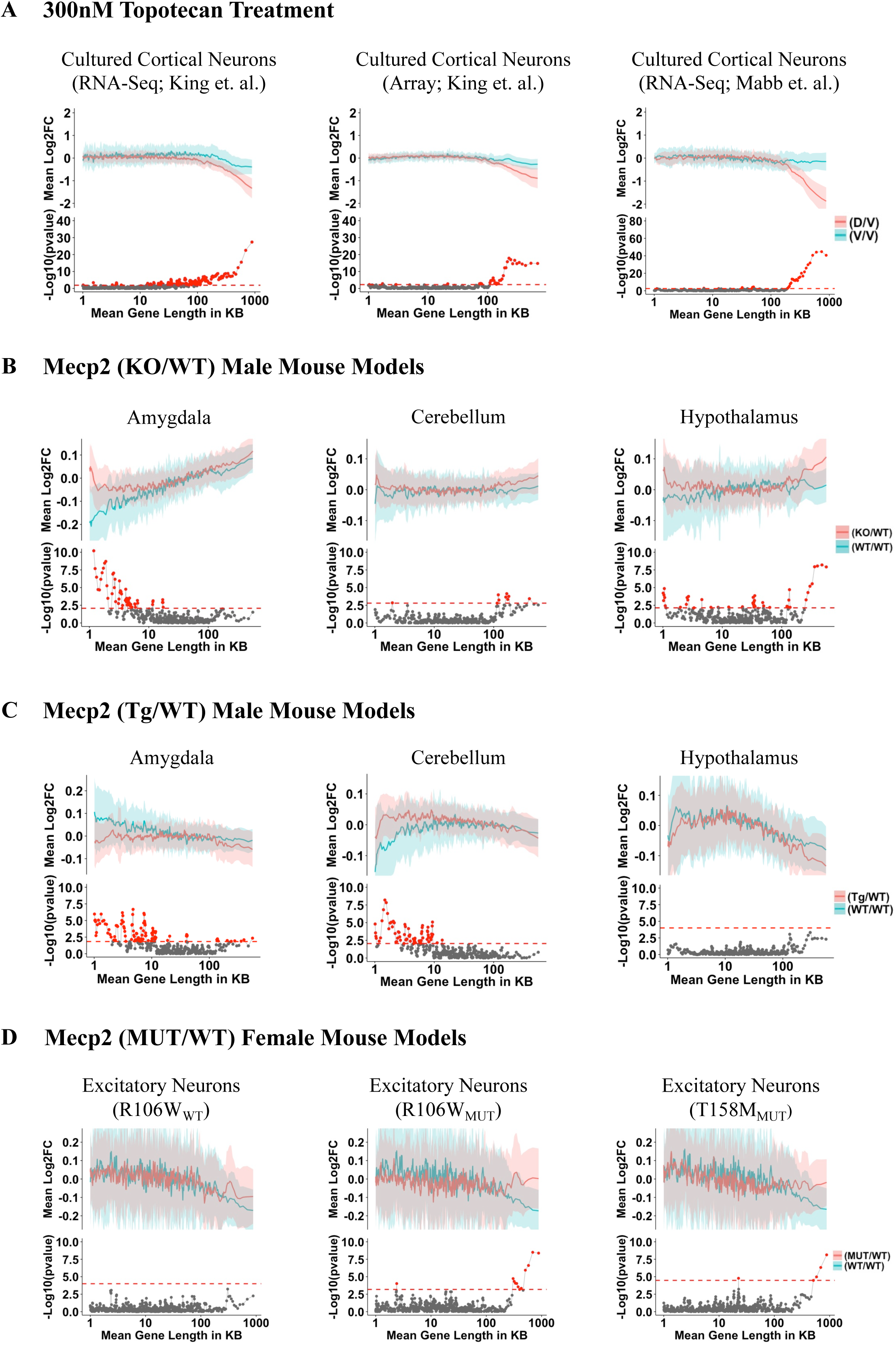
Establishment of baselines and comparison of Mecp2 microarray and RNA-Seq datasets. **(A)** *Topotecan datasets:* The top half of each subgraph shows the comparison of cultured cortical neurons treated with vehicle (V) from C57BL/6J (B6) × CASTEi/J (CAST) F1 hybrid mice with other vehicle-treated samples (V/V, blue line) and comparison of topotecan-treated cortical neurons (D) with vehicle-treated samples (D/V, red line). The red and blue lines diverge only for genes over 100kb in size. (**B-D)** *Mecp2 datasets:* Note the change in the scale of y-axis; these changes are much smaller than in the topotecan studies. Unlike the topotecan results in row A, gene bins with statistical significance are sporadic for both long and short genes in row B. The top half of each subgraph in row **B)** shows the comparison of WT male C57BL samples with other WT male C57BL samples (blue line) and Mecp2 male KO samples compared with WT male littermates (red line) in the amygdala (Samaco et al., 2012), cerebellum (Ben-Shachar et al., 2009) and hypothalamus (Chahrour et al., 2008). **C)** Comparison of three different Mecp2 Tg/WT male mouse models. The top half of each subgraph shows the comparison between WT FVB samples and other WT FVB samples (blue line) within the same genotype and Tg samples with their WT littermates (red line) in amygdala (Samaco et al., 2012), cerebellum (Ben-Shachar et al., 2009) and hypothalamus (Chahrour et al., 2008). Note that we observe few long gene bins as well as short gene bins with significant preferential upregulation in *Mecp2*-null and *Mecp2*-overexpression (Tg) mice datasets. **D)** Cortical excitatory neurons from three different Mecp2 KO/WT female mouse models. The top half of each subgraph shows the comparison between two sets of WT C57BL samples (blue line), and between WT littermates and mutant mice bearing either the R106W or T158M mutations (Johnson et al., 2017). Note that the magnitude of length dependent gene misregulation was more substantial in control samples rather than *Mecp2-mutant* samples (blue curve). The blue or red line represents fold-change in expression for genes binned according to gene length (bin size of 200 genes with shift size of 40 genes) as described in (Gabel et al., 2015). The blue and red shaded areas correspond to one-half of one standard deviation of each bin for the comparison of WT/WT and KO/WT (or MUT/WT) or Tg/WT, respectively. The bottom half of each subgraph is the p-value from the two-sample t-test between KO/WT (or MUT/WT) or Tg/WT and WT/WT. Bins with FDR < 0.05 are shown as a red dot. The red dashed line at the bottom of the subgraphs indicates the minimum -Log_10_(p-value) that corresponds to a FDR < 0.05. Please refer to Table 1 for the total number of samples used for the comparison between two random sets of WT (or vehicle-treated) samples and between WT littermates and KO/Tg/mutant mice.

The first data that we analyzed were those from a study that evaluated transcriptional effects of the topoisomerase 1 inhibitor topotecan in autism (King et al., 2013). When we constructed a running average plot comparing the gene expression changes between topotecan drug-treated neurons (drug or D) and vehicle-treated cortical neurons (vehicle or V), we observed a preferential downregulation of long genes (the running average plot comparing drug vs. vehicle is indicated by the red curve in Figure 1A; Figure S1). To estimate the baseline variation among control samples, two random sets of vehicle-treated cultured cortical neurons were compared to each other (blue curve in Fig 1A, Experimental Procedures). Given that these untreated samples were obtained from littermates, we did not expect to observe any differences in gene expression and predicted that a running average plot comparing gene expression between vehicle-treated control samples would yield a horizontal line through y=0. However, we found that genes over 100kb in length tended to be down-regulated on average (blue curve in Figure 1A) when gene expression levels between control samples are compared. This effect was found for both RNA-Seq and microarray datasets (Figure 1A) and indicates that a portion of the length-dependent trend observed in the topotecan datasets is due to a length-dependent bias (i.e. noise) that can be observed even in the control samples.

To determine the significance of average fold-change trends, we applied a Student’s t-test to each of the matching data bins from the drug vs. vehicle (D/V) and vehicle vs. vehicle (V/V) comparisons, followed by an adjustment for multiple hypothesis testing. For consistency, these plots are referred to as overlap plots (Experimental Procedures, Figure S1). At a false discovery rate of 0.05, only the long gene bins were statistically significant and showed preferential downregulation following topotecan treatment in both RNA-Seq and microarray datasets (lower panel in Fig 1A, red dots indicate statistically significant bins; Figure S1). In other words, although the control samples showed that long genes are downregulated at baseline (i.e. when comparing controls to controls), topotecan treatment produced an even stronger downregulation of long genes, providing sufficient signal to overcome the noise (or intra sample variation) observed in long genes at baseline. These datasets enabled us to establish a statistical procedure as well as provided positive control for further analyses of long gene trends (King et al., 2013; Mabb et al., 2016) in other studies.

### Gene length trends do not hold up in datasets for MeCP2 mouse models

Studies of MeCP2-related disorders—both Rett syndrome (caused by loss-of-function mutations in *MECP2*) and *MECP2* duplication syndrome (caused by duplication or even triplication of the locus)— have provided a wealth of transcriptome data. Experiments in mouse models of both syndromes have suggested that loss of MeCP2 function causes preferential upregulation of long genes and, conversely, that gain of MeCP2 function leads to preferential downregulation of long genes (Gabel et al., 2015). We chose to delve deeper into these datasets to explore the extent of the contribution of long genes to RTT pathology. We applied our method to eleven MeCP2 datasets (Table 2) across seventeen different tissue types (Baker et al., 2013; Ben-Shachar et al., 2009; Chahrour et al., 2008; Chen et al., 2015; Gabel et al., 2015; Kishi et al., 2016; Samaco et al., 2012; Sugino et al., 2014; Zhao et al., 2013). We first computed the running average plots and were able to reproduce the same results as reported previously (Gabel et al., 2015; Sugino et al., 2014). However, when the baseline variation between wild-type (WT) samples is plotted (blue curves in Fig. 1B), they extensively overlap with the running average plots from the *Mecp2*-null (KO) samples (red curves in Fig. 1B; see also Figures S2A-K). This overlap between the curves for the WT vs. WT comparisons and the KO vs. WT comparisons indicates that the signal originally reported for the KO vs. WT comparison can be largely explained by noise (or intra-sample variation) in the dataset, as there is not a clear separation between the WT vs. WT curves and the KO vs. WT curves in most brain regions surveyed.

A few long gene bins showed significant preferential upregulation in *Mecp2*-null mice (FDR < 0.05) in these datasets. For example, in hypothalamus dataset, we found 12 bins of long genes to be significant (Figures 1B, right panel). However, we observed a similar or even larger number of significant bins for genes less than 100k (Figures 1B-1C; see also Figures S2B-S2C, S2F-S2G, and S2J). Likewise, no preferential repression of long genes was observed for datasets from *Mecp2*-overexpression models (Tg) (Figure 1C; see also Figure S2L). Indeed, we found more short genes to be preferentially dysregulated in the *Mecp2*-overexpression models (Figure 1C). Thus, when assessing the bins of genes with the significant difference in expression between WT and KO mice, we found that genes with a variety of lengths were altered in KO and Tg mice. Additionally, while there are certainly some long genes with significantly altered expression in both KO and Tg mice, there is no consistent and preferential long gene trend observed in the *Mecp2* datasets.

### Long gene trend is not present in Nuclear RNA profiles of MeCP2 mouse models

A recent study reported that transcripts of long genes were downregulated in nuclear and nascent RNA samples (Johnson et al., 2017) in contrast to previous studies (Gabel et al., 2015; Sugino et al., 2014). The dataset was generated by combining an *in vivo* biotinylation system with Cre-loxP technology that circumvented cellular heterogeneity of the brain and helped examine transcriptomic changes due to MeCP2 in specific cell types, in both male and female mice (Johnson et al., 2017). The samples were derived from the cortical cells of *Mecp2*-mutant mice bearing either of two common Rett-causing mutations: T158M or R106W, which are among the most common mutations found in RTT patients (Cuddapah et al., 2014).

We reanalyzed the data using overlap plots and observed no significant downregulation of long genes in wildtype or *Mecp2*-mutant excitatory neurons from 18-week old T158M or R106W female mice (Figures 1D and S4E). In excitatory neurons bearing the R106W mutation, we observed few bins that are significantly different from WT expression levels. Notably, bins with significant gene expression changes were not due to the downregulation of long genes in mutant samples. Rather, these bins were significant due to the downregulation of long genes in control (WT) samples, as indicated by the downward slopes of the running average plots comparing WT vs. WT samples (blue lines in Figures 1D and S4E). Similarly, we observed no significant repression of long genes in nuclear RNA-Seq datasets of excitatory and inhibitory neurons from 6-week old male mice with the same mutation type (Figures S4A-S4D). Finally, when we examined downregulation of long genes from the GRO-Seq (global nuclear run-on with high-throughput sequencing) data collected from these mice, we confirmed a marginal significance in the downregulation of long genes (Figure S3A), but upregulation of long genes was not observed in whole cell RNA-Seq data (Figure S3B). These results suggest that the transcriptome changes in long genes that appear in RNA isolation-based methods are independent of the sex, age, or mutation type of the mouse.

Together, these results suggest that when the fold-change difference is 50% or more, as it is in the topotecan datasets, there is likely to be a genuine long gene bias. When the fold-change effect is small (<15%), however, as it is with the long genes observed in the *Mecp2* datasets, it is more likely that the observed long gene trend is due to inherent variation among samples. The reported long gene trend in the *Mecp2* datasets is in the same range as the noise that we derived from the intra-sample comparison in the control groups, and this effect was seen in all the *Mecp2* datasets that we assessed. This further suggests that the length-dependent variability estimated from microarray and RNA-Seq platforms is not sensitive enough to capture small transcriptional changes. We, therefore, recommend that baseline gene length dependency should be evaluated from the control group first to understand the statistical significance of observed long gene trends in any sequencing dataset.

### Human MeCP2 datasets: the importance of age

To determine whether preferential dysregulation of long genes occurs in *in vitro* human Rett datasets, we computed overlap plots on samples from isogenic human iPSCs (hiPSCs), neural progenitor cells (NPCs), and neurons from the fibroblasts of two independent patients, with and without the *MECP2* mutation. We found no preferential upregulation of long genes (Figures 2A) but did see a trend toward downregulation of long genes among human *in vitro* RTT neuron samples, which is contrary to reports from *Mecp2*-null mouse models (Gabel et al., 2015; Sugino et al., 2014).

**Figure 2.**
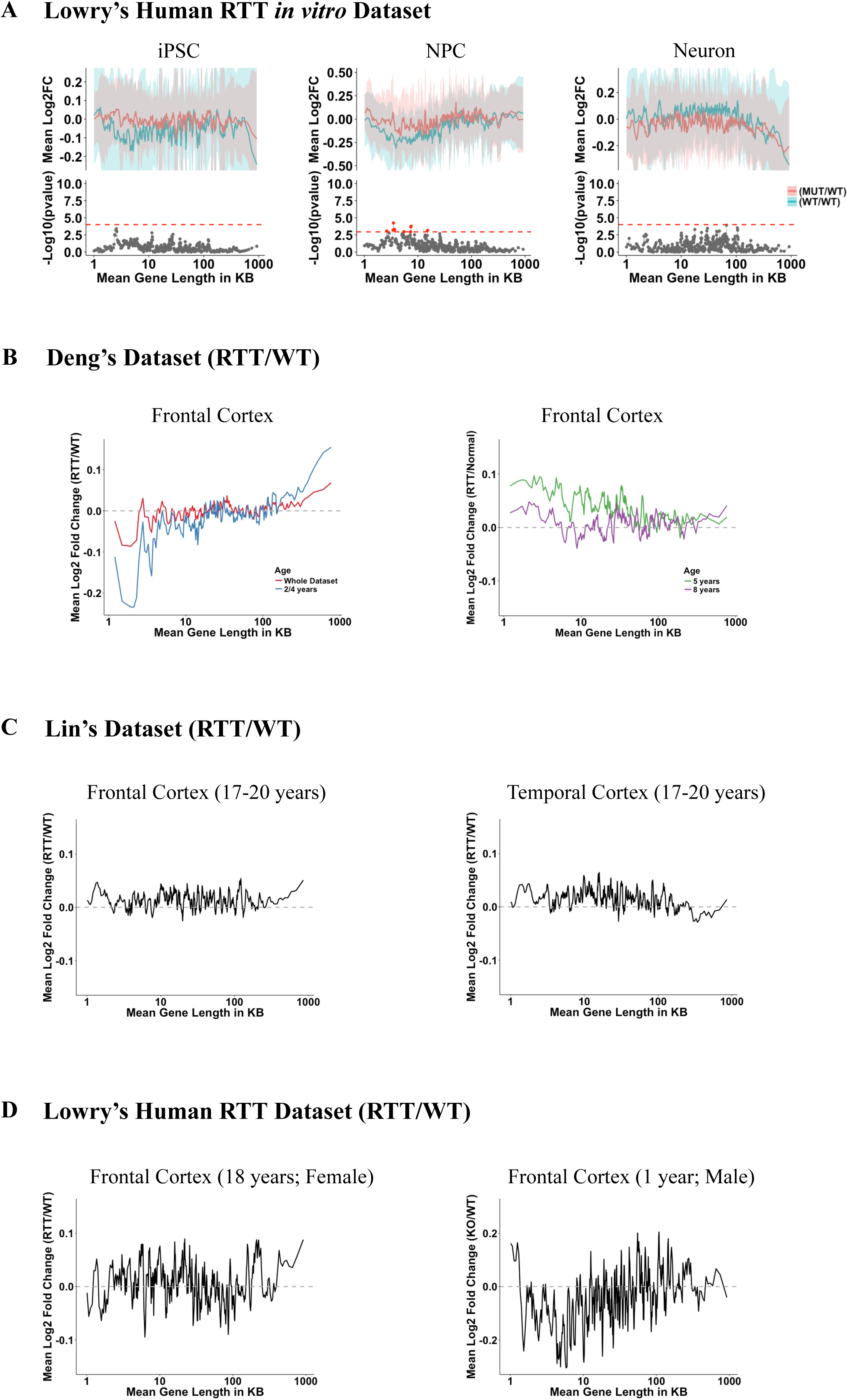
No bias toward long genes in *MECP2* human datasets. **(A)** RNA-Seq analysis of isogenic human Rett *in vitro* models. Overlap plots were used to compare WT and KO samples, where the top half of each subgraph shows the comparison of WT samples with other WT samples (blue line), and RTT samples compared with WT samples (red line) in iPSC (left panel), Neural progenitor cells or NPC (middle panel), and neurons (right panel). **(B)** Microarray analysis of human RTT brain samples compared to age-matched control for Frontal Cortex (Deng et al., 2007). Comparison of gene trends in the pooled sample from 2-and 4-year old patients (left panel) and whole dataset (left panel). Observed long gene trend in the sample from 5-year old (right panel) and 8-year old patients (right panel). **(C)** Microarray analysis of RTT human frontal cortex samples (Lin et al., 2016) compared to controls (left panel) and RTT human temporal cortex samples (Lin et al., 2016) compared to controls (right panel). **(D)** RNA-Seq analysis of RTT human (female) frontal cortex samples compared to controls (left panel) and RTT human (male) frontal lobe samples compared to controls (right panel). The lines in A-D represent fold-change in expression for genes binned according to gene length (bin size of 200 genes with shift size of 40 genes) as described in Gabel, Kinde et al. *Nature* 2015. The blue and red ribbons in **(A)** correspond to one-half of one standard deviation of each bin for the comparison of WT/WT and MUT/WT respectively. The bottom half of each subgraph is the p-value from the two-sample t-test between MUT/WT and WT/WT. Bins with FDR < 0.05 are shown as a red dot. The red dotted line in the bottom of the subgraphs indicates the minimum -Log_10_(p-value) that corresponds to a FDR < 0.05. Please refer to Table 1 for the total number of samples used for the comparison between two random sets of WT samples and between WT and RTT samples.

Although long genes do not appear to be upregulated above the level of background noise in murine *Mecp2* datasets, they have been reported to be preferentially upregulated in human RTT samples (Gabel et al., 2015) as well, and we wondered if a more robust signal would be observed in post-mortem human datasets. Three RTT and three normal control samples from the superior frontal gyrus were obtained from a previous study (Deng et al., 2007). These samples were from three different ages: RTT samples were obtained from donors aged 8, 6, and <4 years (pooled samples from a 2- and a 4-year old), with approximately age-matched normal control samples obtained from donors aged 10, 5 and 2 years, respectively. The long gene trend was observed (Gabel et al., 2015) in a comparison of the three RTT samples to the three control samples (Figures 2B). Because stages of brain development and disease progression in RTT patients change markedly from ages 1 to 5 years before stabilizing (Chahrour and Zoghbi, 2007), we reanalyzed the data by comparing each sample to its age-matched control separately. Dysregulation of long genes was observed only in the 2- and 4-year old RTT samples (Figure 2B left panel), but not in either the 5- or 8-year old RTT samples (Figure 2B right panel). Unfortunately, the statistical significance of this observation cannot be established because of the small sample size (n = 1 each).

To determine whether length-dependent misregulation of long genes occurs in other human datasets, we analyzed samples from another study (Lin et al., 2016) and in-house generated RNA-Seq RTT datasets. Lin et al. dataset (Lin et al., 2016) consist of postmortem brain samples from the frontal and temporal cortex of RTT patients with age-matched controls (age = 17–20 years, n = 3 each). Because the phenotypes are similar for RTT patients in this age range (Chahrour and Zoghbi, 2007), we grouped these RTT samples together and compared them to the pooled age-matched controls. We computed running average plots on the normalized dataset (Experimental Procedures, Figure S1) and did not observe overrepresentation of long genes in these samples (Figure 2C). Similar results were reported by the original study (Lin et al., 2016). Consistent with our previous results, there was no long gene trend in the running average plot of the RNA-Seq RTT dataset collected from a postmortem frontal cortex sample obtained from an 18-year-old RTT female (Figure 2D, left panel) when it was compared to its age-matched control (age = 18 years, n = 1 each). To further probe whether the long-gene trend might be present in the early stages of the disease, we compared a RTT postmortem male sample from frontal cortex (age = 1 year, n = 1) to an age-matched control sample (age = 2 days, n = 1) and again could find no significant upregulation of long genes (Figure 2D, right panel).

One possible explanation for the lack of a long gene trend in human RTT samples is heterogeneity among the various samples (including differences in the genetic background), which increases the inherent variability in gene expression among biological replicates. Such variability could obscure the effects of a subtle bias in the sequencing process. Nevertheless, the present findings suggest that long genes are not preferentially misregulated in human RTT datasets.

### Differential gene expression analysis for Topotecan and Mecp2 datasets

Our previous analyses suggest that the current transcriptome profiling technologies are limited in their ability to detect subtle differences in gene expression. We hypothesize that long gene effects, if genuine, should be apparent in both binning analysis and the traditional differential gene expression analysis. We, therefore, decided to focus our attention on only the differentially expressed genes that were reported by previous studies (Baker et al., 2013; Ben-Shachar et al., 2009; Chahrour et al., 2008; Chen et al., 2015; Huang et al., 2011; King et al., 2013; Mabb et al., 2016). We divided the entire list of differentially expressed genes into four groups based on gene length (> or < 100kb) and fold-change direction (either up or down). Consistent with our overlap plots, we found long genes to be substantially overrepresented and downregulated in Topotecan datasets (Figure 3A). This result proves that our approach does detect long gene trends in gene expression studies. In the MeCP2 datasets, however, we did not find a preferential upregulation of long genes (Figures 3B-3D) except in the hippocampal dataset (Figure S5) (Baker et al., 2013). Moreover, in contrast to previous studies, we found that more long genes were upregulated than downregulated in the cerebellum of *Mecp2* over-expressing mice (Figure 3C, right panel). Another important difference between the Topotecan and *Mecp2* datasets was that short genes dominated among all differentially expressed genes in *Mecp2* datasets (Figures 3B-3D; Figures S5). This further supports the notion that a preference for long gene misregulation is not an inherent feature of gene expression following the *Mecp2* disruption. This is not to say that MeCP2 does not regulate a subset of long genes, only that our analysis found no preferential misregulation of long gene trend in MeCP2 mouse models.

**Figure 3.**
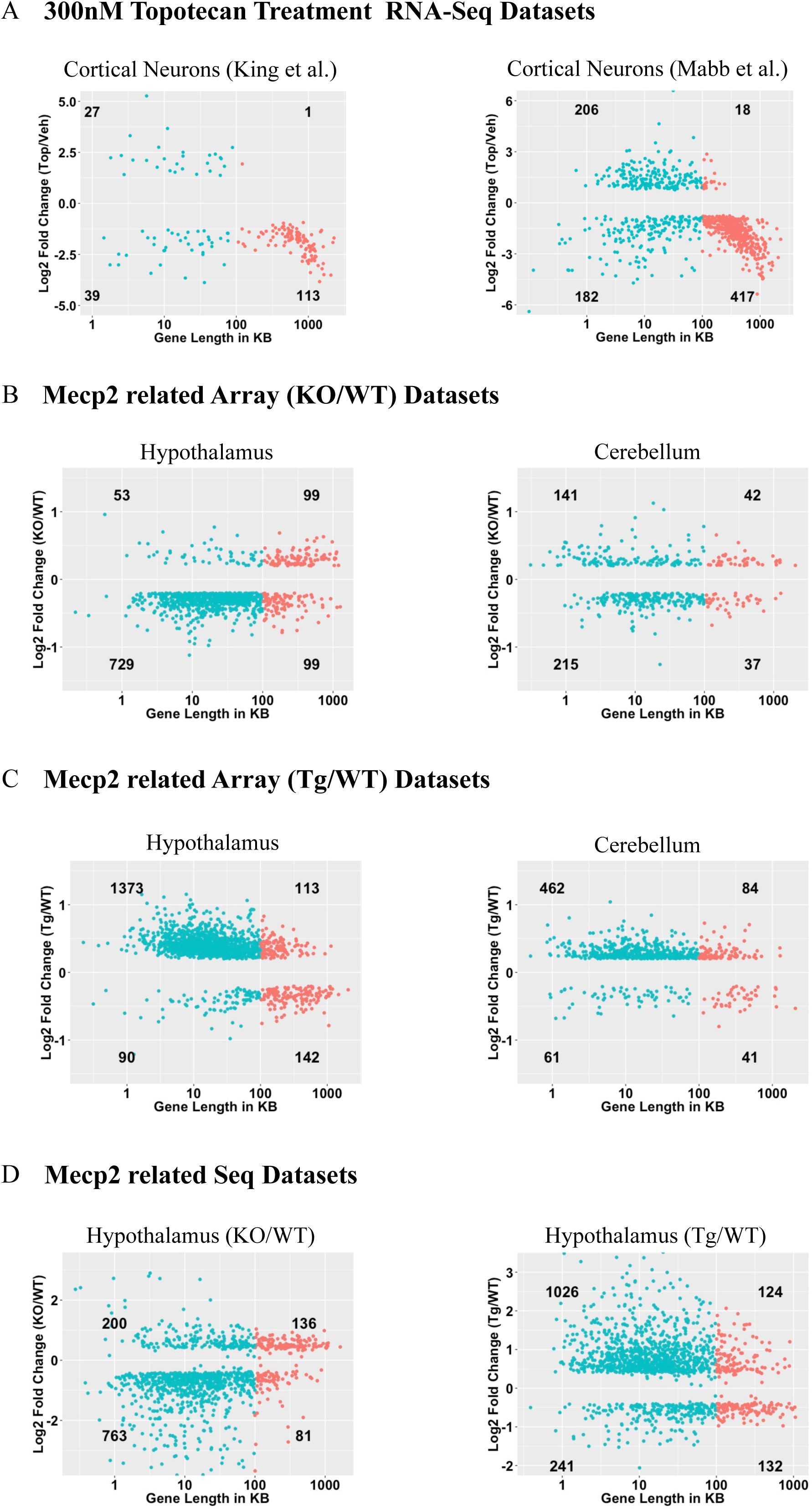
Differentially expressed genes show length-dependent misregulation in Topotecan datasets but not in Mecp2 studies. **(A)** Scatter plot of log fold-change in expression between topotecan and vehicle-treated cultured cortical neurons (y-axis) against its gene length (x-axis) in RNA-Seq dataset from (King et al., 2013) (left panel; n = 5 each; FDR < 0.05) and (Mabb et al., 2016) RNA-Seq dataset (right panel; n = 3 each; FDR < 0.01). **(B)** Scatter plot of log fold-change in expression (microarray) between C57BL KO and its C57BL WT littermates (y-axis) against its gene length (x-axis) in hypothalamus (left panel; n = 4 each; FDR < 0.05 and log2FC > 0.2; (Chahrour et al., 2008)) and cerebellum (right panel; n = 4 each; FDR < 0.05 and log2FC > 0.2; (Ben-Shachar et al., 2009)). **(C)** Scatter plot of log fold-change in expression (microarray) between FVB Tg to its FVB WT littermates (y-axis) against its gene length (x-axis) in hypothalamus (n = 4 each; FDR < 0.05 and log2FC > 0.2; (Chahrour et al., 2008)) and cerebellum (n = 4 each; FDR < 0.05 and log2FC > 0.2; (Ben-Shachar et al., 2009)). **(D)** Scatter-plot of log fold-change in expression between KO/Tg and WT littermates (y-axis) against gene length (x-axis) in RNA-Seq datasets: Hypothalamus KO/WT comparison (left panel; n = 3 each; FDR < 1e-5; (Chen et al., 2015)) and Hypothalamus Tg/WT comparison (right panel; n = 3 each; FDR < 1e-5; (Chen et al., 2015)). Red dot represents long genes and blue dot represents short genes. Differentially expressed genes were obtained from the published gene lists.

### RNA-Seq and microarray benchmark datasets are prone to length-dependent bias

To investigate whether length dependent bias might be a function of amplification-based platforms, we next performed running average analysis on the samples from the phase-III Sequencing/Microarray Quality Control (SEQC) project (Consortium, 2014). SEQC was designed to evaluate the performance of various sequencing platforms, sources of bias in gene expression samples, and various methods for downstream analysis. The consortium generated benchmark datasets using four different types of RNA samples: A (Universal Human Reference RNA), B (Human Brain Reference RNA), C (a mixture of A and B at a ratio of 3:1), and D (a mixture of A and B at a defined ratio of 1:3). The RNA-Seq datasets generated using the Illumina HiSeq 2000 platform across six different sites were used for quality control analyses (Experimental Procedures), and the raw read counts were normalized using the DESeq2 method (Love et al., 2014).

To determine whether the dataset showed nominal batch effects or other non-biological variability, we used multidimensional scaling (MDS) plots to see if the samples clustered according to RNA sample type. To ascertain whether or not the samples were consistently titrated, we calculated the β ratio of observed gene expression in the samples, which is obtained from the following equation: ((B-A)/(C-A)) (Consortium, 2014). The value of the β ratio (Shippy et al., 2006) is 4:1 (or log2(4) = 2). In theory, the β ratio should be independent of gene length in the brain and non-brain tissues. After assessing various SEQC datasets, we found that the Novartis dataset had nominal batch effects and the β ratio was close to 2. Therefore, this dataset would be ideal, as it would not bias downstream analyses (Figure 4A).

**Figure 4.**
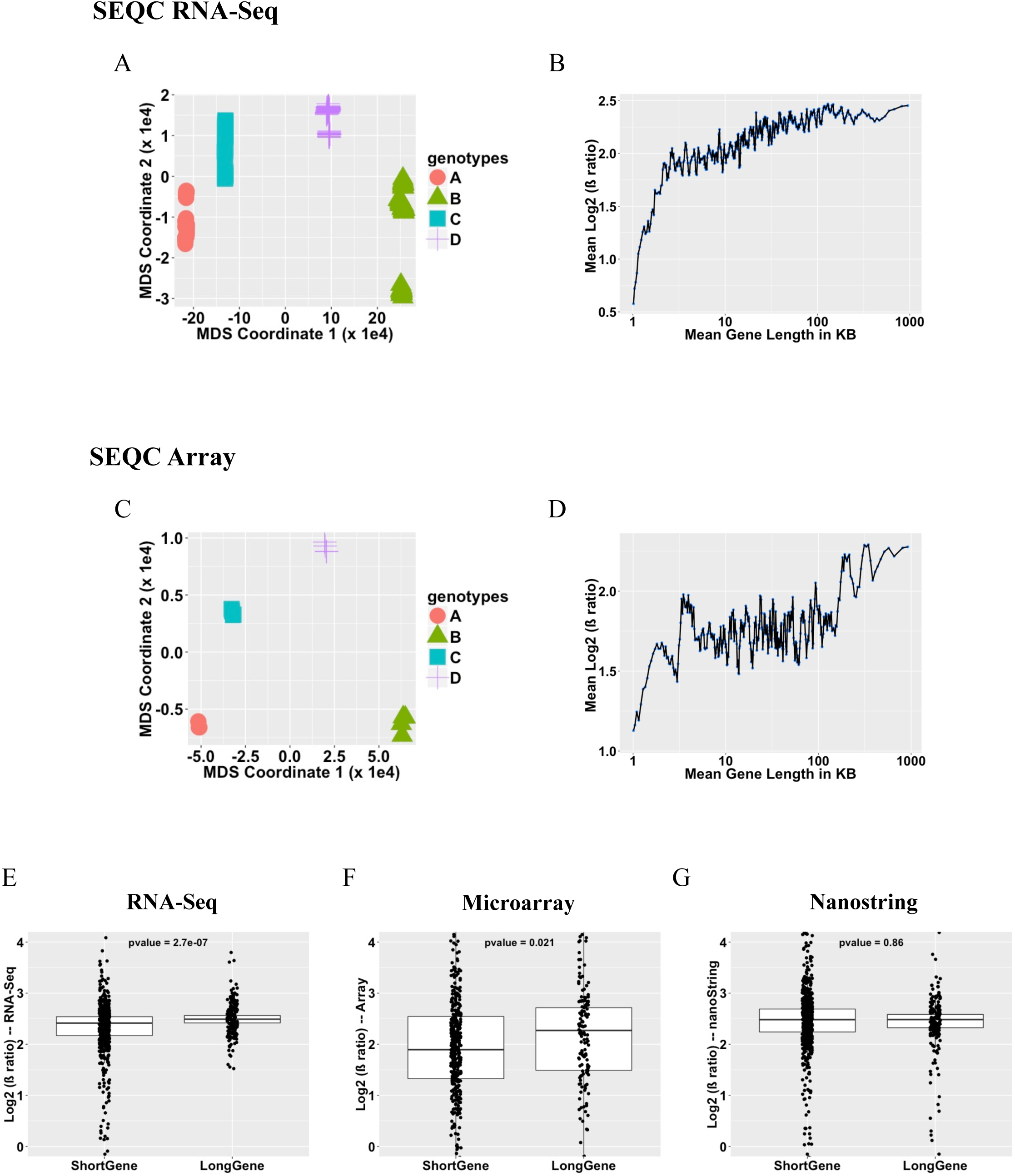
Long gene bias in SEQC RNA-Seq and microarray, but not NanoString, datasets. **(A)** MDS plot using Euclidean distance on the SEQC (Consortium, 2014) NVS count dataset. **(B)** Mean Log2 Fold Change plot against gene length using β ratio samples ((B-A/C-A); n =64 each) in RNA-Seq dataset. **(C)** MDS plot using Euclidean distance on the SEQC microarray dataset. **(D)** Mean Log2 Fold-Change plot against gene length using β ratio samples in microarray dataset (n = 4 each). Each blue dot is a bin of 200 genes with shift size of 40 genes (Gabel et al., 2015). Box plot of the genes across three different platforms that are present in NanoString codeset. The distributions of the mean fold-changes for β ratio samples for long and short genes are compared across three different platforms: **E)** RNA-Seq, **F)** Microarray, and **G)** NanoString. P-values were computed using the Wilcoxon Mann Whitney test.

We separated Human Brain Reference (sample type B) RNA-Seq samples into two groups of 32 samples each, based on their y-axis coordinates of the MDS plot, and computed a running average plot. Since these samples were technical replicates of the same reference RNA sample type, we expected the mean log2 fold-change to be a horizontal line along the x-axis with a y-intercept equal to zero (i.e., y=0 on an xy plane). Instead, we found that long genes deviated from the expected pattern, with the fold-changes of long genes being overestimated (Figure S6A, left panel).

We then investigated whether the fold-change of long genes is constant for the P ratio samples. The expected average log2 fold-change should be a horizontal line along the x-axis with a y-intercept equal to two. We found, however, that the expected ratio was not maintained for long genes and was overestimated (Figure 4B). Moreover, we observed a similar bias in the β ratio with respect to transcript length, with longer transcripts being overrepresented (Figure S6A, right panel). Overall, the range of overestimation in the RNA-Seq dataset was between 3% and 40%. Consistent with our findings, another study (using a different dataset) previously reported that long genes were more likely to be identified as statistically significant in RNA-Seq datasets (Oshlack and Wakefield, 2009).

To determine whether the long gene bias was unique to the RNA-Seq datasets or could be detected on other platforms, we investigated the MAQC-III microarray Affymetrix dataset generated by the SEQC consortium (Consortium, 2014). Human Brain Reference samples (B) were separated into two groups based on y-axis location on the MDS plot (Figure 4C). The running average plots were computed against their average gene length using the same parameters as described for the RNA-Seq analysis above. As with the RNA-Seq samples, the average fold-change for long genes deviated from the expected value of zero (Figure S6B, left panel). When the β ratio was plotted against the mean gene length (Figure 4D) or mean transcript length (Figure S6B right panel), we found that long genes were overrepresented. Further, long gene bias was observed in both RNA-Seq and microarray datasets in a comparison of two groups of universal human reference (Figure S6A-S6B, middle panel). The overestimation in the microarray dataset ranged from 1.5% to 23%—lower overall than for the RNA-Seq dataset, but indicating that microarray datasets are also predisposed to gene and transcript length-dependent bias.

### Long gene bias is independent of normalization methods

To ensure that the long gene bias we observed was not due to our normalization methods, we compared the mean log2 fold-change using three different normalization techniques: Total Count, DESeq (Anders and Huber, 2010), and edgeR/TMM (Robinson et al., 2010; Robinson and Oshlack, 2010). We normalized the raw read counts from four different RNA sample types using each of the three normalization methods and computed running average plots of the P ratios against gene and transcript length. In all cases, long genes were still overestimated, regardless of the normalization method (Figures S7A-S7B). This lends support to the notion that the overrepresentation of long genes is independent of the normalization technique.

### Long gene bias is not observed in NanoString datasets, which are not based on amplification

We hypothesized that PCR amplification, a process shared by both microarray and RNA-Seq technologies, might introduce the observed bias in long gene expression. We, therefore, performed NanoString nCounter gene expression quantification, a technique that does not use amplification, with the SEQC reference RNA samples (A, B, C, and D) (n = 6 each). The MDS plot on normalized data showed that the samples clustered based on sample type (Figure S8A); the effect of batches was minimal (Experimental Procedures). The code set consisted of ~ 184 long genes, out of which ~132 long genes were expressed in brain samples (Figure S8B). We again computed the running average plots against their average gene length, and we did not observe any long gene bias between the brain samples or when computing the β ratio of the samples (Figures S8C-S8D).

We next compared the mean expression levels of all the common genes across the RNA-Seq, microarray and nCounter datasets. Our analysis shows that fold-changes of long genes are overestimated in the RNA-Seq (P-value < 2.7 e-07; Figure 4E) and microarray datasets (P-value < 0.021; Figure 4F); in contrast, the nCounter dataset showed no difference in the average expression of long and short genes (P-value = 0.86; Figure 4G). Although it is possible that the smaller number of genes (~680) might make it more difficult to detect a preference, the proportion of long genes in this dataset (~180 out of ~680 genes, or 26%) is twice that found in the human transcriptome (~3200 long genes out of ~ 24,000 genes, or 13%). Any preference for long genes should thus be revealed even more strongly in this dataset. These results lead us to posit that the long gene overestimation we observed in RNA-Seq and microarray datasets might be caused by a length-dependent bias in PCR amplification.

### PCA plot confirms the reciprocal relationship of Mecp2 gain- and loss-of-function datasets

One of the most intriguing components of the long gene story in RTT is the presence of a reciprocal pattern in the *Mecp2*-overexpression model, where a reported preference for downregulation of long genes complements the upregulation of long genes reported in *Mecp2*-null mice (Gabel et al., 2015). To understand this reciprocal relationship, we divided Human Brain Reference samples (B) into 3 groups (n = 16 each) based on different library preparation ID numbers from the Novartis SEQC dataset. The PCA plot clearly clustered the brain samples based on the library preparation group to which they belonged (Figure S9A). Comparing the brain samples of library preparation ID 2 (green) to library preparation ID 1 (red) and ID 3 (blue) separately reversed the running average plot (Figures S9B-S9C). These results show that a reciprocal relationship can be observed in the gene expression data between any groups that form three distinct clusters on a PCA plot.

We next assessed the influence of the fold-change threshold on differential expression analysis using brain samples. Although we did not expect to see a trend between replicates, preferential regulation of long genes was observed (Figure S9D) when the fold-change was small (<10%, or log2FC ~ 13%).

The bias was similar to the trend observed in previously published *Mecp2*-null and overexpression (Tg) models when library preparations ID 2 (red) and ID 1 (green), or library preparations ID 3 (blue) and ID 1 (green), were compared (Gabel et al., 2015; Sugino et al., 2014).

In this analysis, all the samples were technical replicates of the same reference RNA and were expected to have identical gene expression levels, but variation associated with library preparation resulted in the samples not clustering together and allowed us to observe an inverse trend in long genes (Figure S9A). Just as biological variation can lead to separation on a PCA plot, so can technical variation, and both can result in the same apparent long gene bias observed in *Mecp2* datasets. Furthermore, our analysis suggests that differentially expressed genes can be highly variable with small fold-changes, which underscores the importance of proper fold-change cut-offs in differential gene expression analysis.

### Differentially expressed genes with small fold-changes identified by RNA-Seq are not reproducible by NanoString in the *Mecp2* dataset

To determine whether a long gene trend is present only in the *Mecp2* RNA-Seq dataset and not in the NanoString dataset, we generated RNA-Seq (> 90 million paired-end sequencing reads per sample; n = 3 each; Table 3) and NanoString (n = 3 each; Table 4) datasets on cerebellar tissue from wild-type and *Mecp2*-null mouse models (KO). The PCA plot on normalized datasets (Experimental Procedures) showed that the samples clustered based on sample type (Figures S10A-S10B, left panel). Transcriptome analysis was performed using DESeq2 (Love et al., 2014) on both datasets. We first analyzed RNA-Seq data to estimate the strength of the long gene trend. Although there appeared to be a long gene trend in the KO/WT comparison, an overlap plot confirmed there was no significant upregulation of long genes (Figure S10A, middle panel). Consistent with our previous findings, there was no preferential upregulation of long genes in our differential expression analysis (Figure S10A, right panel; absolute log2FC > 1.2 & FDR < 0.05).

We performed further analysis using a list of 750 (~159 long and ~591 short) genes common to both RNA-Seq and nCounter NanoString (Experimental Procedures). Comparison of the log fold-changes using the classic method (i.e., log2((mean(group1) + 1)/(mean(group2) + 1)) and using shrunken log fold-changes by DESeq2 (i.e., obtaining reliable variance estimates by pooling information across all the genes) suggested that the latter method yields more highly correlated fold-changes (Figures S10C). This is consistent with previous findings showing that shrunken log fold-changes are more reproducible (Love et al., 2014; Robinson et al., 2010). Even with this method, however, we observed high variability among genes with low fold-changes between the two datasets, regardless of whether they were long or short (Figures 5A and 5B). Moreover, genes with high fold-changes in expression (~ FC > 20%) were consistently called as differentially expressed in both the datasets (Figures 5A and 5B).

**Figure 5.**
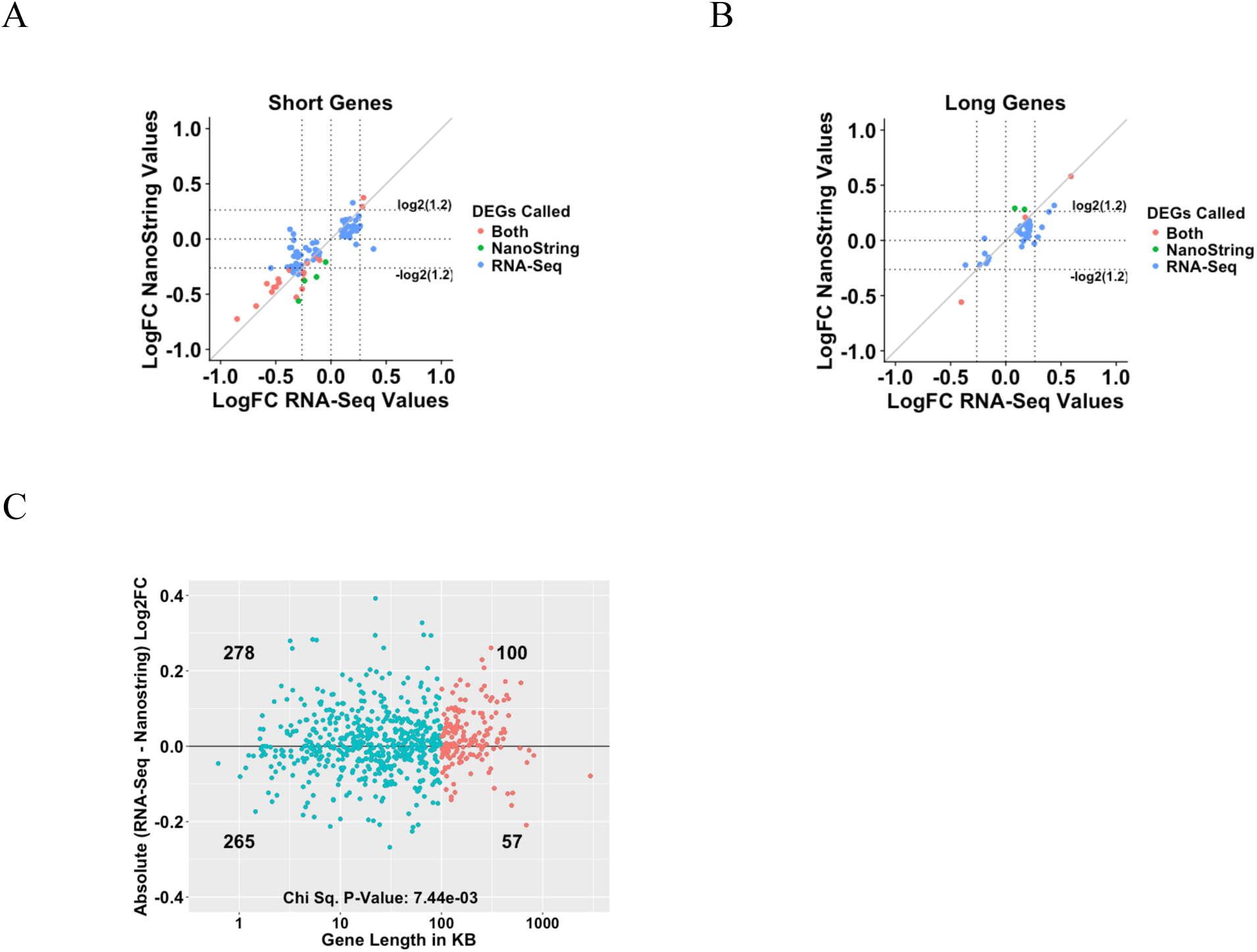
Expression changes are overestimated in RNA-Seq datasets. Comparison of log fold-change in expression between RNA-Seq and Nanostring for Short Genes (**A**) and Long Genes (**B**). Here, we used FDR < 0.05 for a gene to be considered differentially expressed. A Red dot represents genes that are called as differentially expressed by both platforms. The Green and Blue dot represents genes that are called differentially expressed by Nanostring and RNA-Seq respectively. **C)** Absolute log fold-change difference between RNA-Seq and Nanostring (y-axis) against gene length (x-axis). A Red dot represents long gene and blue dot represents short genes. P-values were computed using chi-square test.

**Table 1:** List of Comparisons used in overlap or average plots

| Brain region | Mouse strain/Human samples compared | Reference |
| --- | --- | --- |
| **Fig 1A.** Cultured Cortical Neurons (left panel) | BL: Hybrid Vehicle vs Hybrid Vehicle (n = 2 each)<br>RL: Hybrid Topotecan vs Hybrid Vehicle (n = 5 each) | King et al. *Nature* 2013 |
| Cultured Cortical Neurons (middle panel) | BL: Hybrid Vehicle vs Hybrid Vehicle (n = 1 each)<br>RL: Hybrid Topotecan vs Hybrid Vehicle (n = 3 each) | King et al. *Nature* 2013 |
| Cultured Cortical Neurons (right panel) | BL: Hybrid Vehicle vs Hybrid Vehicle (n = 1 each)<br>RL: Hybrid Topotecan vs Hybrid Vehicle (n = 3 each) | Mabb et al. *PLoS One* 2016 |
| **Fig 1B.** Amygdala (left panel) | BL: C57BL WT vs C57BL WT (n = 2 each)<br>RL: C57BL KO vs C57BL WT (n = 5 each) | Samaco et al. *Nature Genetics* 2012 |
| Cerebellum (middle panel) | BL: C57BL WT vs C57BL/6J WT (n = 2 each)<br>RL: C57BL KO vs C57BL/6J WT (n = 5 each) | Ben-Shachar et al., *Human Mol. Genet.* 2009 |
| Hypothalamus (right panel) | BL: C57BL WT vs C57BL/6J WT (n = 2 each)<br>RL: C57BL KO vs C57BL/6J WT (n = 4 each) | Chahrour et al., *Science* 2008 |
| **Fig 1C.** Amygdala (left panel) | BL: FVB WT vs FVB WT (n = 2 each)<br>RL: FVB KO vs FVB WT (n = 5 each) | Samaco et al. *Nature Genetics* 2012 |
| Cerebellum (middle panel) | BL: FVB WT vs FVB WT (n = 2 each)<br>RL: FVB KO vs FVB WT (n = 5 each) | Ben-Shachar et al. *Human Mol. Genet.* 2009 |
| Hypothalamus (right panel) | BL: FVB WT vs FVB WT (n = 2 each)<br>RL: FVB KO vs FVB WT (n = 4 each) | Chahrour et al., *Science* 2008 |
| **Fig 1D.** Cortical Excitatory Neurons R106W<sub>WT</sub> (left panel) | BL: C57BL WT vs C57BL WT (n = 1 each)<br>RL: C57BL R106W<sub>WT</sub> vs C57BL WT (n = 2 each) | Johnson et al., *Nature Medicine* 2017 |
| Cortical Excitatory Neurons R106W<sub>MUT</sub> (middle panel) | BL: C57BL WT vs C57BL WT (n = 1 each)<br>RL: C57BL R106W<sub>MUT</sub> vs C57BL WT (n = 2 each) | Johnson et al., *Nature Medicine* 2017 |
| Cortical Excitatory Neurons T158M<sub>MUT</sub> (right panel) | BL: C57BL WT vs C57BL WT (n = 1 each)<br>RL: C57BL T158M<sub>MUT</sub> vs C57BL WT (n = 2 each) | Johnson et al., *Nature Medicine* 2017 |
| **Fig 2A.** iPSC (left panel) | BL: iPSC WT vs iPSC WT (n = 2 each)<br>RL: iPSC RTT (n = 4) vs iPSC WT (n = 5 each) | GSE# |
| NPC (middle panel) | BL: NPC WT vs NPC WT (n = 2 each)<br>RL: NPC RTT (n = 4) vs NPC WT (n = 5 each) | GSE# |
| Neuron (right panel) | BL: Neuron WT vs Neuron WT (n = 2 each)<br>RL: Neuron RTT vs Neuron WT (n = 4 each) | GSE# |
| **Fig 2B.** Frontal Cortex (left panel) | RL: Post mortem RTT vs Controls (n = 3 each)<br>BL: Post mortem pooled sample from 2- and 4-year old patient vs Control (n = 1 each) | Deng et al. *Human Mol. Genet.* 2007 |
| Frontal Cortex (right panel) | GL: Post mortem pooled sample from 5-year old patient vs age matched control (n = 1 each)<br>PL: Post mortem pooled sample from 8-year old patient vs age matched control (n = 1 each) | Deng et al. *Human Mol. Genet.* 2007 |
| **Fig 2C.** Frontal Cortex (left panel) | RTT female samples compared to age matched controls (ages 17-20 years; n = 3) | Lin et al. *BMC Genomics* 2016 |
| Temporal Cortex (right panel) | RTT female samples compared to age matched controls (ages 17-20 years; n = 3) | Lin et al. *BMC Genomics* 2016 |
| **Fig 2D.** Frontal Cortex (left panel) | RTT female samples compared to age matched controls (ages 18 years; n = 1 each) | GSE# |
| Frontal Cortex (right panel) | RTT male samples (age 1 year) to compared to age matched (age 2 day) controls (n = 1 each) | GSE# |

Note: Hybrid mouse line is C57BL/6J (B6) × CAST/Ei/J (CAST) F1 hybrid mice. BL, RL, GL and PL stands for Blue line, Red line, Green line and Purple line respectively.

This analysis suggests that the genes identified as differentially expressed by RNA-Seq at lower fold changes are not reproducible by NanoString. To determine whether fold-changes are inflated in RNA-Seq, we compared the absolute difference of log2 fold-change between the RNA-Seq and NanoString datasets. We observed fold-changes of long genes to be overestimated by RNA-Seq technology (Figure 5C; Chi-Square test; p-value <7.44e-3), which further supports our hypothesis that artefactual long gene trends are more likely to appear in amplification-based expression datasets.

## DISCUSSION

Several recent papers have suggested that diseases associated with synaptic dysfunction tend to preferentially involve misregulation of long genes (>100 Kb) (Gabel et al., 2015; King et al., 2013; Sugino et al., 2014; Zylka et al., 2015). To establish a statistical baseline for the length-dependent gene regulation analysis, we took advantage of a large number of SEQC consortium datasets where the relative gene expression fold-change has been measured using RNA-Seq and microarray. We demonstrated the power of big data analysis by uncovering major sources of technical variation such as intra-sample variation and PCR amplification bias that can affect the analysis of long gene expression. By contrast, NanoString nCounter technology, which does not rely on amplification, revealed no long gene bias. Our results demonstrate that amplification-based transcriptomic technologies can lead to overestimations of long gene expression changes.

This is not to say that there is never a bias toward expression changes in long genes. The topotecan dataset showed an authentic long gene trend even after accounting for baseline variability. This sizeable effect on long gene expression is consistent with the biological function of topotecan inhibiting topoisomerase I; long genes should, in theory, be more dependent on proper unwinding during transcription elongation (King et al., 2013). By contrast, we found no bias toward long gene dysregulation in the MeCP2 datasets after baseline correction, even when we focused on only those genes that are differentially expressed to a statistically significant degree. The sole exception was the one infantile RTT case, but a single case does not allow us to draw any firm conclusions. Again, this does not rule out that MeCP2 regulates some long genes; it simply does not support a preferential misregulation of long genes by mutant MeCP2.

Apparent expression changes in long genes are clearly liable to exaggeration by biases in microarray and RNA-Seq. We recommend eliminating confounds such as batch effects and properly estimating both inter- and intra-sample variations; the control datasets must be carefully analyzed in order to reveal the degree of baseline variability, which then can inform further analyses of the size of the signal required to overcome background noise in sequencing datasets (Figure S1). These findings are applicable to all research that utilizes current microarray and sequencing technologies. We hope that revealing the influence of protocols and technologies on RNA sequencing data will lead to improved technologies and more reliable analyses for amplification-based sequencing data.

## AUTHOR CONTRIBUTIONS

Conceptualization: ZL, HYZ, ATR, AEP; Methodology and Investigation: ATR, ZL; Software, Validation and Analysis: ATR, YWW, AEP; Data Curation: ATR, BL, AEP, YWW, HKY; Writing – Original Draft: ATR, ZL, AEP; Writing – Review & Editing: ATR, ZL, AEP, YWW; Visualization: ATR, YWW; Supervision: ZL, HYZ; Funding Acquisition: ZL, HYZ.

## ACKNOWLEDGMENTS

We thank Laura Lavery, Rami Al-Ouran, Laura Lombardi, Ezequiel Sztainberg, Aya Ishida, and Vicky Brandt for helpful discussions and suggestions. This project was supported by the Genomic and RNA Profiling Core at Baylor College of Medicine and the expert assistance of the Core Director, Lisa D. White, Ph.D.

## ACCESSION NUMBERS

The GEO accession numbers for NanoString and RNA-seq datasets reported in this paper are as follows: GSE94073, GSE105047 (includes GSE105045 and GSE105046) and GSE107399.

## Supplementary Figure Legends

**Figure S1.**
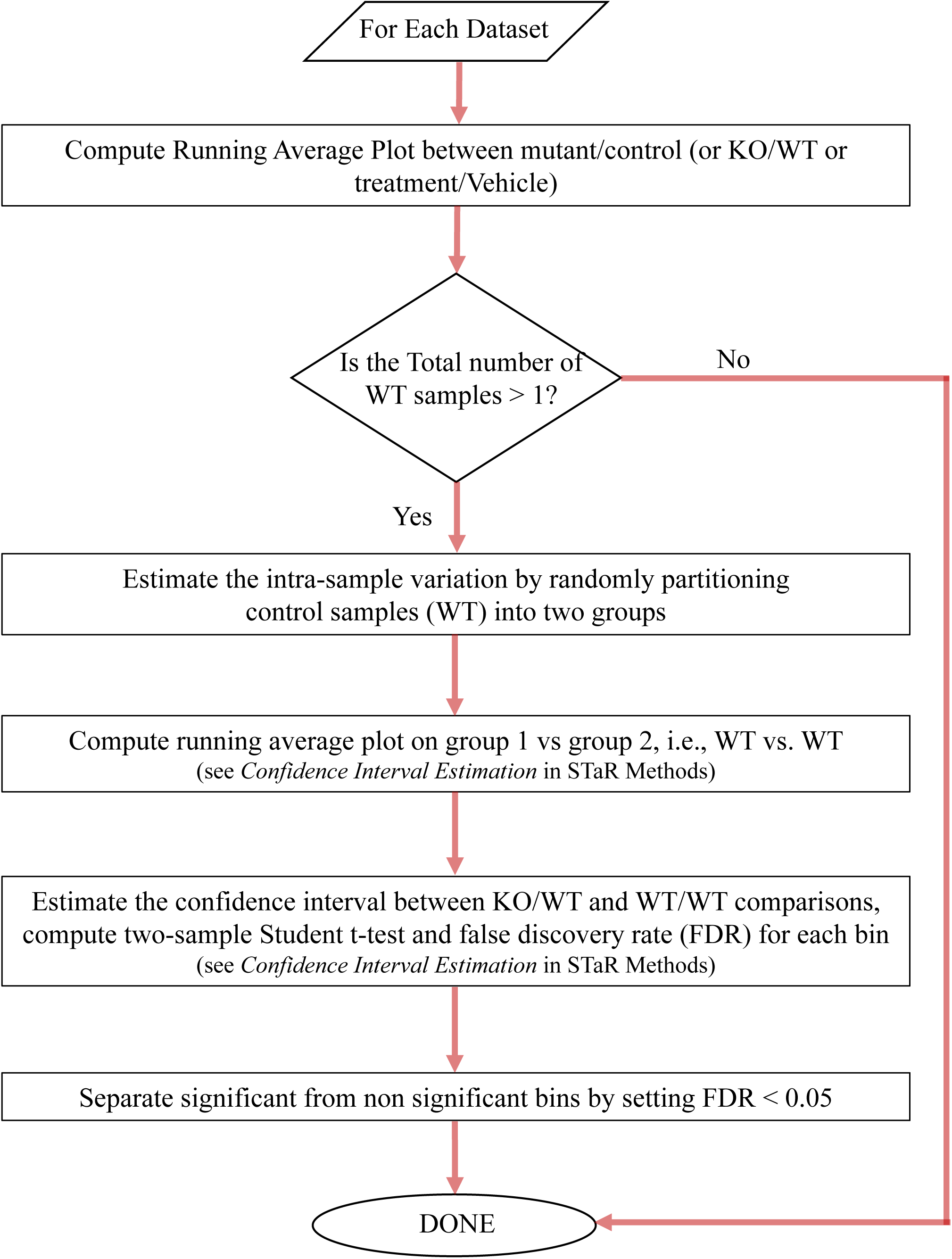
Schematic diagram of rigorous assessment of long gene trends (Related to Figure 1).

**Figure S2.**
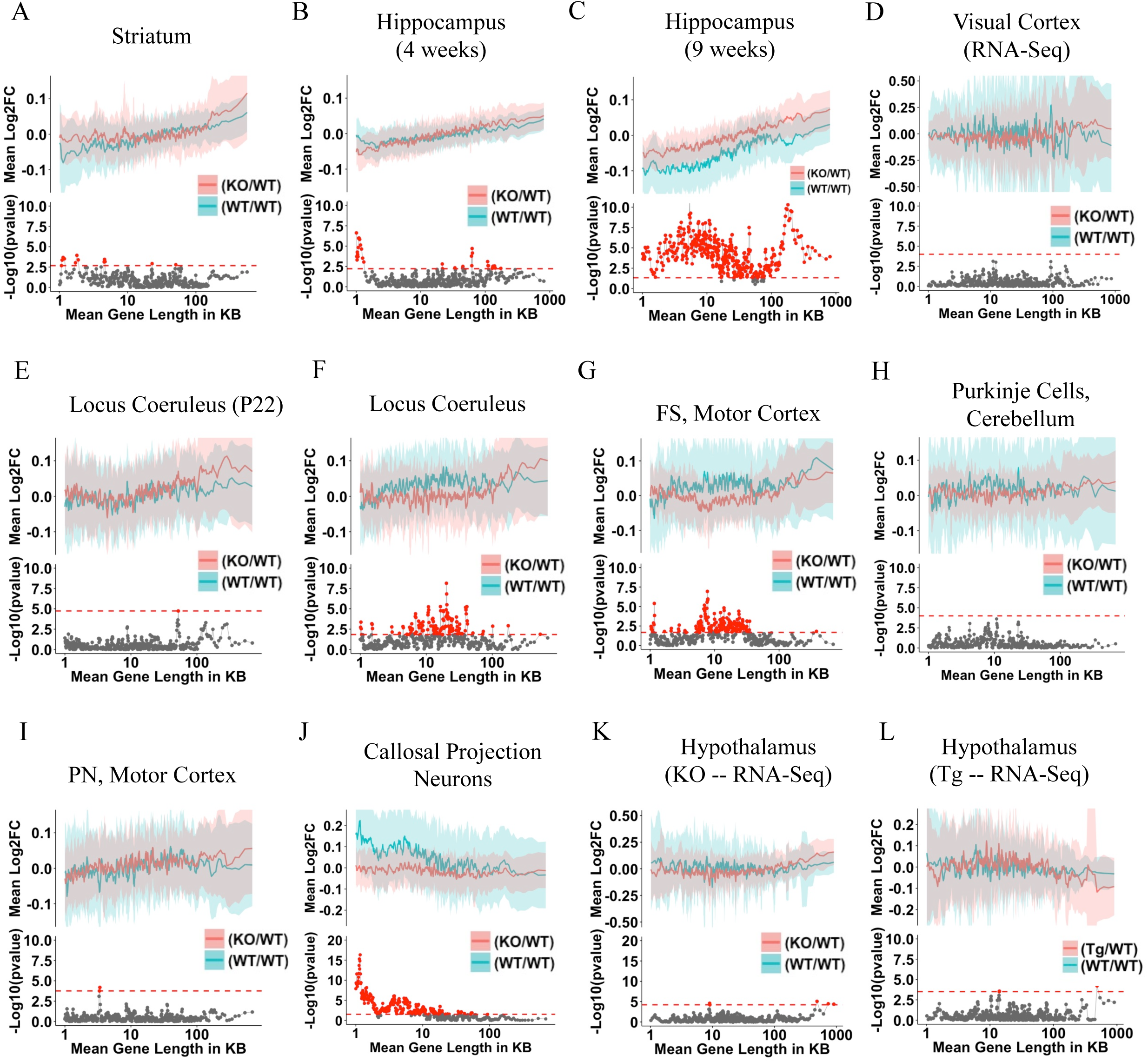
Long gene trend is not present in Mecp2 datasets (Related to Figure 1). (A-L) Analysis of Intra-sample variation in WT *Mecp2* dataset shows a bias toward long genes across different brain regions. The Blue line (BL) represents the comparison of permuted WT/WT samples from a respective dataset (as mentioned in the comparison table). The Red line (RL) represents the comparison of KO/MUT/Tg samples to its WT littermates from a respective dataset (as mentioned in the comparison table). The top half of each subgraph shows the lines that represent fold-change in expression for genes binned according to gene length (bin size of 200 genes with shift size of 40 genes) as described (Gabel et al., 2015; Zhao et al., 2013). Note that we observe few long gene bins as well as short gene bins with significant preferential upregulation in *Mecp2*-null mice datasets. The blue and red ribbon correspond to one-half of one standard deviation of each bin for the comparison of WT/WT and KO/WT or Tg/WT, respectively. The bottom half of each subgraph is the p-value from the two-sample t-test between KO/WT or Tg/WT and WT/WT. Bins with FDR < 0.05 are showed in red. The red dotted line indicates the minimum -Log_10_(p-value) that corresponds to a FDR < 0.05. Here is the list of comparisons for Figure S2:

| Brain region | Mouse lines compared | Reference |
| --- | --- | --- |
| A. Striatum | BL: C57BL WT vs C57BL WT (n = 2 each)<br>RL: C57BL KO vs C57BL WT (n = 5 each) | Zhao et al. *Neuro of Disease* 2013 |
| B. Hippocampus (4 weeks) | BL: FVBx129 WT vs FVBx129 WT (n = 2 each)<br>RL: FVBx129 KO vs FVBx129 WT (n = 4 each) | Baker et al. *Cell* 2013 |
| C. Hippocampus (9 weeks) | BL: FVBx129 WT vs FVBx129 WT (n = 2 each)<br>RL: FVBx129 KO vs FVBx129 WT (n = 4 each) | Baker et al. *Cell* 2013 |
| D. Visual Cortex | BL: WT vs WT (n = 1 each)<br>RL: KO (Mecp2tm1.1Bird) vs WT (n = 3 each) | Gabel, Kinde et al. *Nature* 2015 |
| E. Locus Coeruleus Neurons (TH Young/~P22) | BL: C57BL/6J WT vs C57BL/6J WT (n = 1 each)<br>RL: C57BL/6J KO vs C57BL/6J WT (n = 3 each) | Sugino et al. *J Neurosci.* 2014 |
| F. Locus Coeruleus Neurons (TH) | BL: C57BL/6J WT vs C57BL/6J WT (n = 1 each)<br>RL: C57BL/6J KO vs C57BL/6J WT (n = 3 each) | Sugino et al. *J Neurosci.* 2014 |
| G. Fast Spiking interneurons, Motor Cortex (G42) | BL: C57BL/6J WT vs C57BL/6J WT (n = 2 each)<br>RL: C57BL/6J KO vs C57BL/6J WT (n = 4 each) | Sugino et al. *J Neurosci.* 2014 |
| H. Purkinje Cells, Cerebellum (G42) | BL: C57BL/6J WT vs C57BL/6J WT (n = 1 each)<br>RL: C57BL/6J KO vs C57BL/6J WT (n = 3 each) | Sugino et al. *J Neurosci.* 2014 |
| I. Pyramidal Neurons, Motor Cortex (YPFH) | BL: C57BL/6J WT vs C57BL/6J WT (n = 1 each)<br>RL: C57BL/6J KO vs C57BL/6J WT (n = 3 each) | Sugino et al. *J Neurosci.* 2014 |
| J. Callosal Projection Neurons | BL: C57BL/6J WT vs C57BL/6J WT (n = 1 each)<br>RL: C57BL/6J KO vs C57BL/6J WT (n = 3 each) | Kishi et al. *Nature Comm.* 2016 |
| K. Hypothalamus (KO – RNA-Seq) | BL: FVBx129SvEvTac WT vs FVBx129SvEvTac WT (n=1 each)<br>RL: FVBx129SvEvTac KO vs FVBx129SvEvTac WT (n = 3 each) | Chen et al. *PNAS* 2015 |
| L. Hypothalamus (Tg – RNA-Seq) | BL: FVBx129SvEvTac WT vs FVBx129SvEvTac WT (n=1 each)<br>RL: FVBx129SvEvTac Tg vs FVBx129SvEvTac WT (n = 3 each) | Chen et al. *PNAS* 2015 |

**Figure S3.**
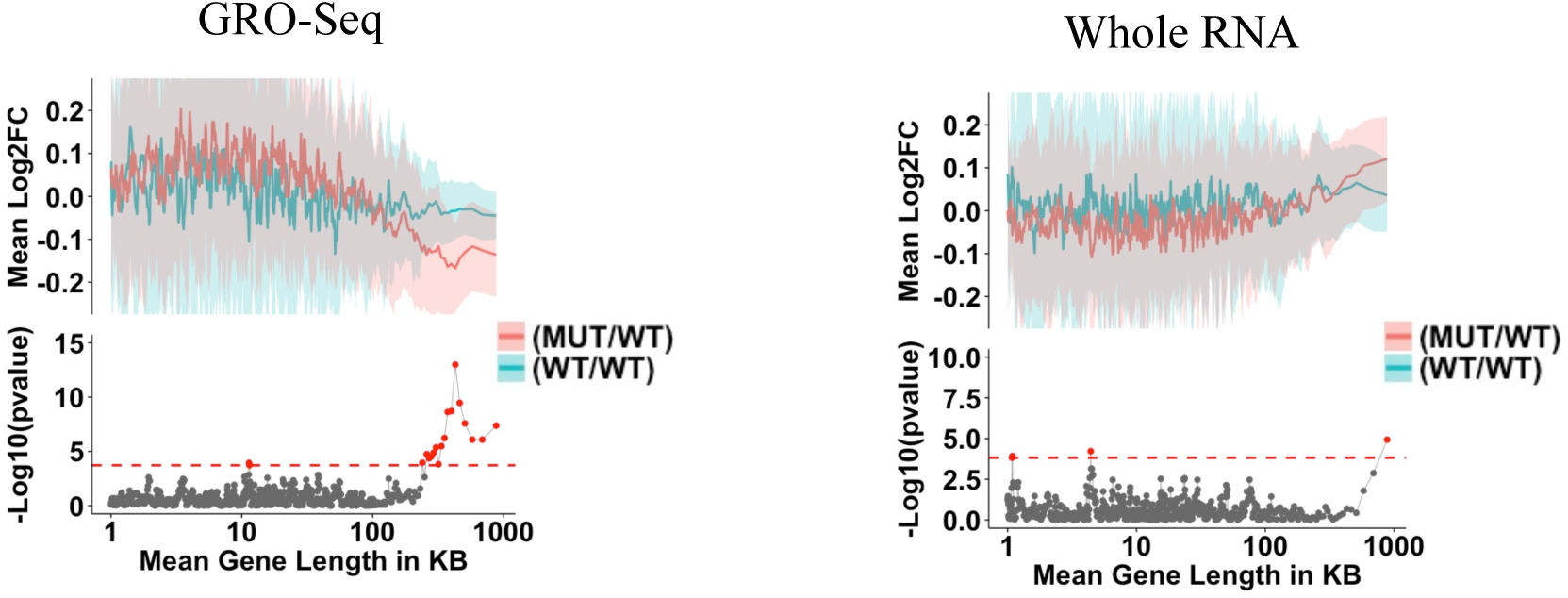
Intra-sample variation bias in WT *Mecp2* datasets is independent of the RNA isolation method (Related to Figure 1): Blue line (BL) represents the comparison of permuted WT/WT samples from a respective dataset (as mentioned in the comparison table). The Red line (RL) represents the comparison of MUT samples to its WT littermates from a respective dataset (as mentioned in the comparison table). The top half of each subgraph shows the lines that represent fold-change in expression for genes binned according to gene length (bin size of 200 genes with shift size of 40 genes) as described (Gabel et al., 2015). The blue and red ribbon correspond to one-half of one standard deviation of each bin for the comparison of WT/WT and MUT/WT respectively. The bottom half of each subgraph is the p-value from the two-sample t-test between MUT/WT and WT/WT. Bins with FDR < 0.05 are shown in red. The red dotted line indicates the minimum -Log_10_(p-value) that corresponds to a FDR < 0.05.

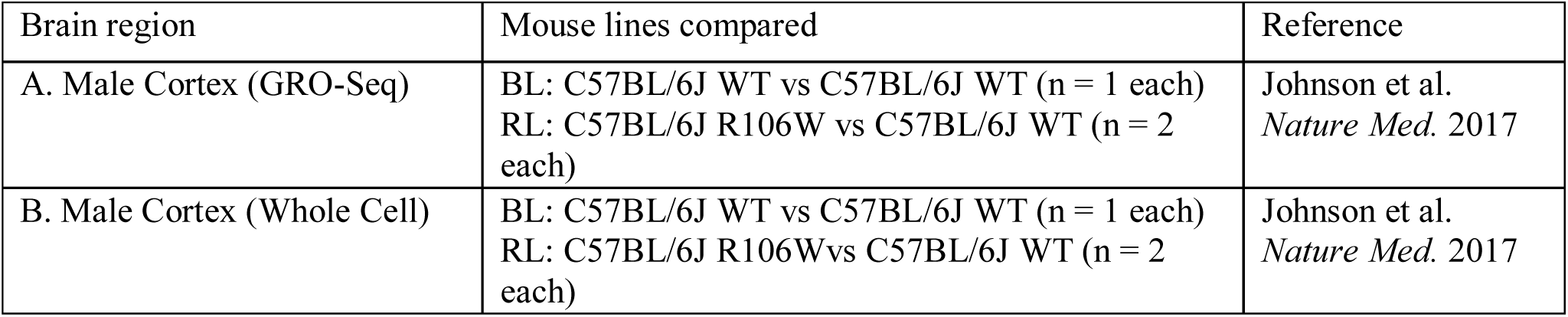

| Brain region | Mouse lines compared | Reference |
| --- | --- | --- |
| A. Male Cortex (GRO-Seq) | BL: C57BL/6J WT vs C57BL/6J WT (n = 1 each)<br>RL: C57BL/6J R106W vs C57BL/6J WT (n = 2 each) | Johnson et al.<br>*Nature Med.* 2017 |
| B. Male Cortex (Whole Cell) | BL: C57BL/6J WT vs C57BL/6J WT (n = 1 each)<br>RL: C57BL/6J R106W vs C57BL/6J WT (n = 2 each) | Johnson et al.<br>*Nature Med.* 2017 |

**Figures S4.**
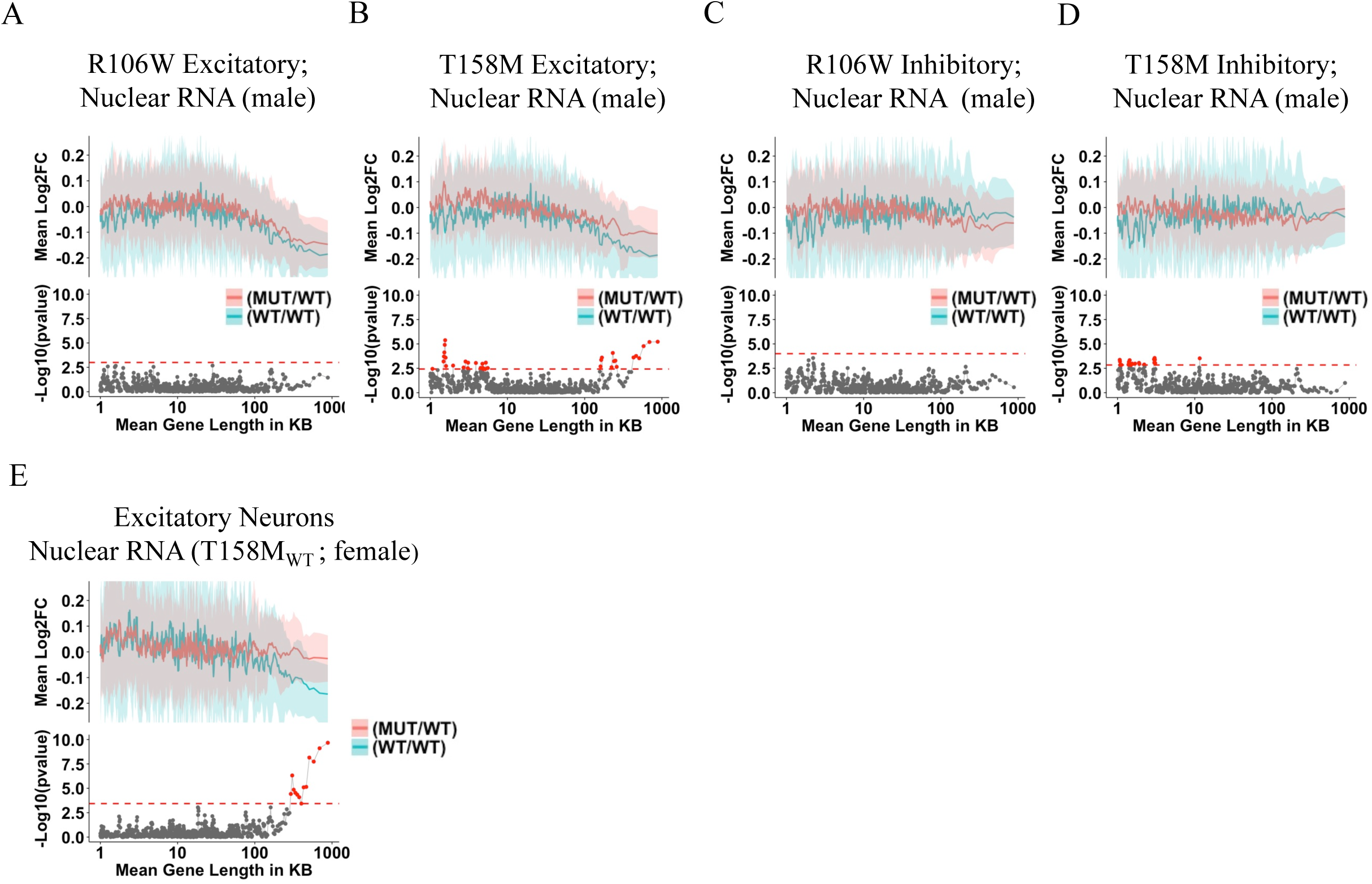
Intra sample variation bias in WT *Mecp2* datasets is independent of the sex of mouse of the mouse model (Related to Figure 1): Blue line (BL) represents the comparison of permuted WT/WT samples from a respective dataset (as mentioned in the comparison table). The Red line (RL) represents the comparison of MUT samples to its WT littermates from a respective dataset (as mentioned in the comparison table). The top half of each subgraph shows the lines that represent fold-change in expression for genes binned according to gene length (bin size of 200 genes with shift size of 40 genes) as described (Gabel et al., 2015). The blue and red ribbon correspond to one-half of one standard deviation of each bin for the comparison of WT/WT and MUT/WT respectively. The bottom half of each subgraph is the p-value from the two-sample t-test between MUT/WT and WT/WT. Bins with FDR < 0.05 are shown in red. The red dotted line indicates the minimum -Log_10_(p-value) that corresponds to a FDR < 0.05.

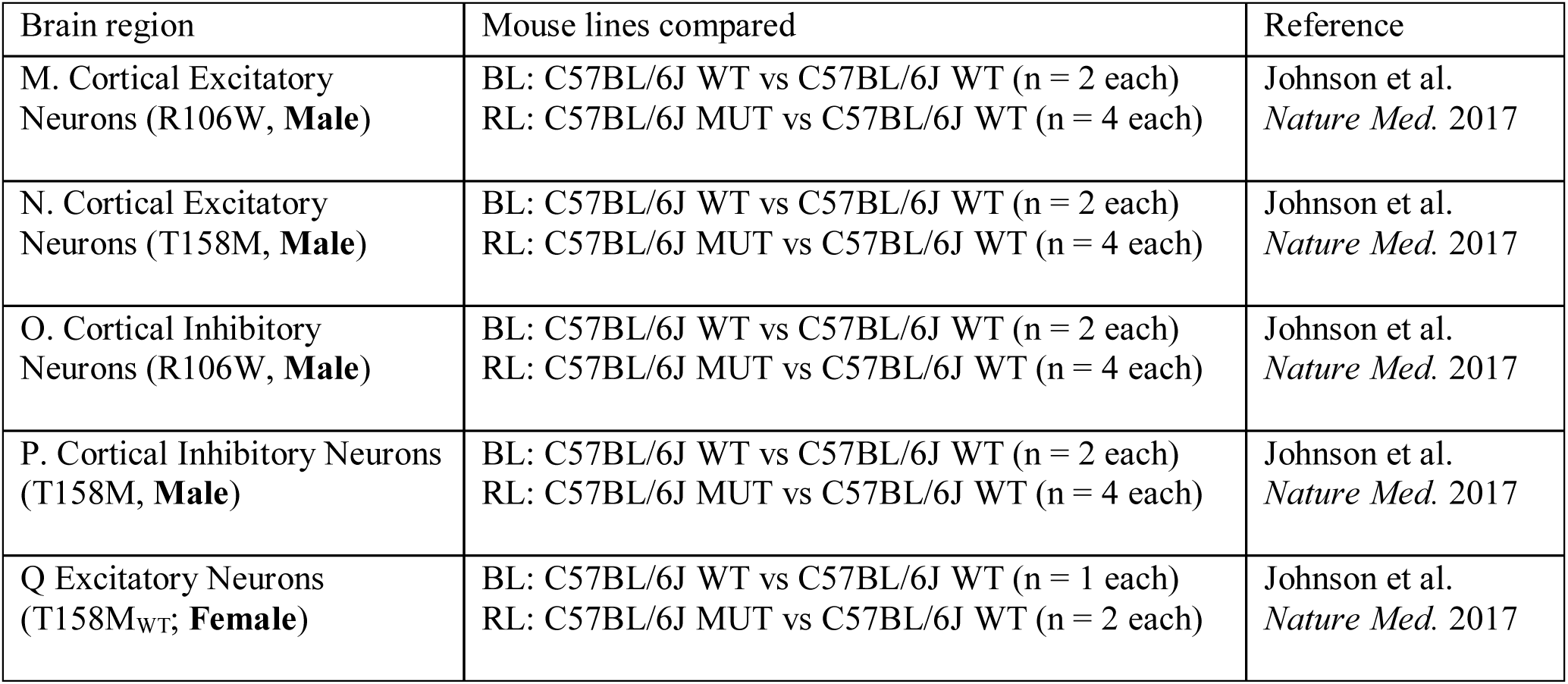

**Figure S5.**
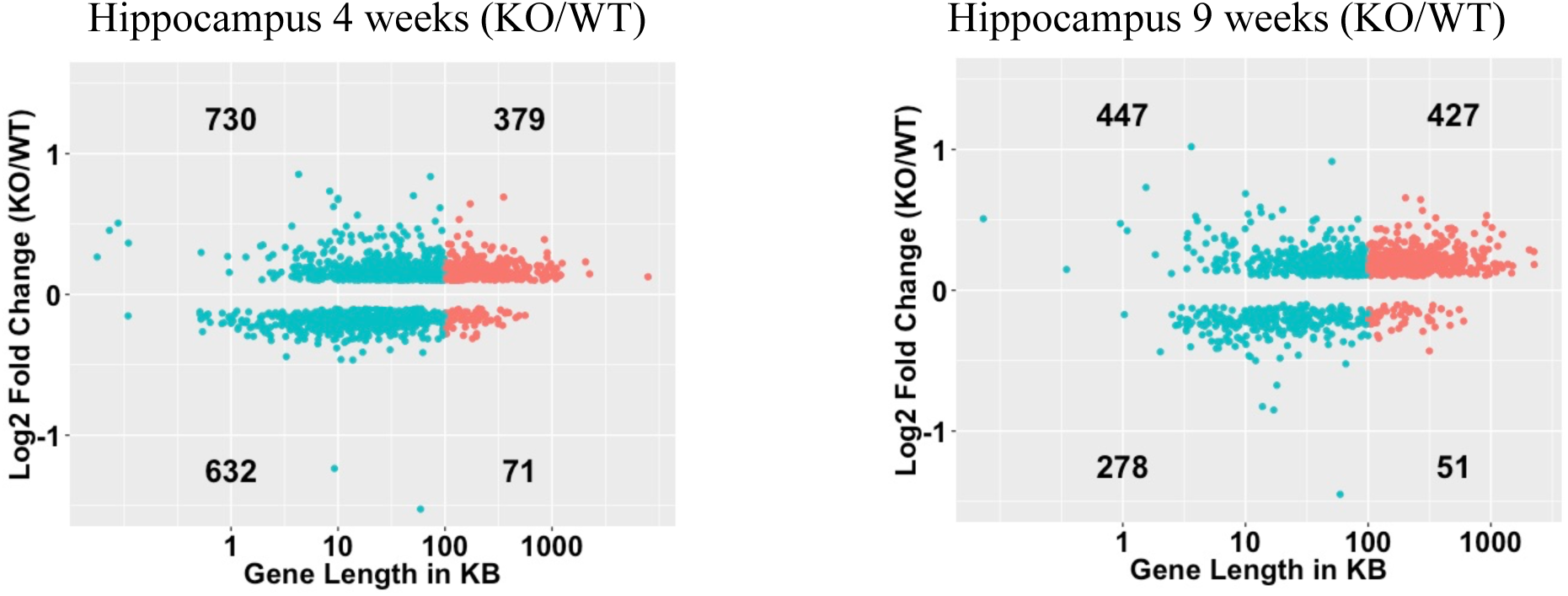
(Related to Figure 3). Differentially expressed gene analysis using gene list from Baker et al., 2013 on Mecp2 hippocampus dataset. Scatter plot of log fold-change (log2FC > 0.1 and FDR < 0.05) in expression between FVBx129 KO to its FVBx129 WT littermates (y-axis) against its gene length (x-axis) in samples of hippocampus from 4-week old and 9-week old mice (n = 4; (Baker et al., 2013)).

**Figure S6.**
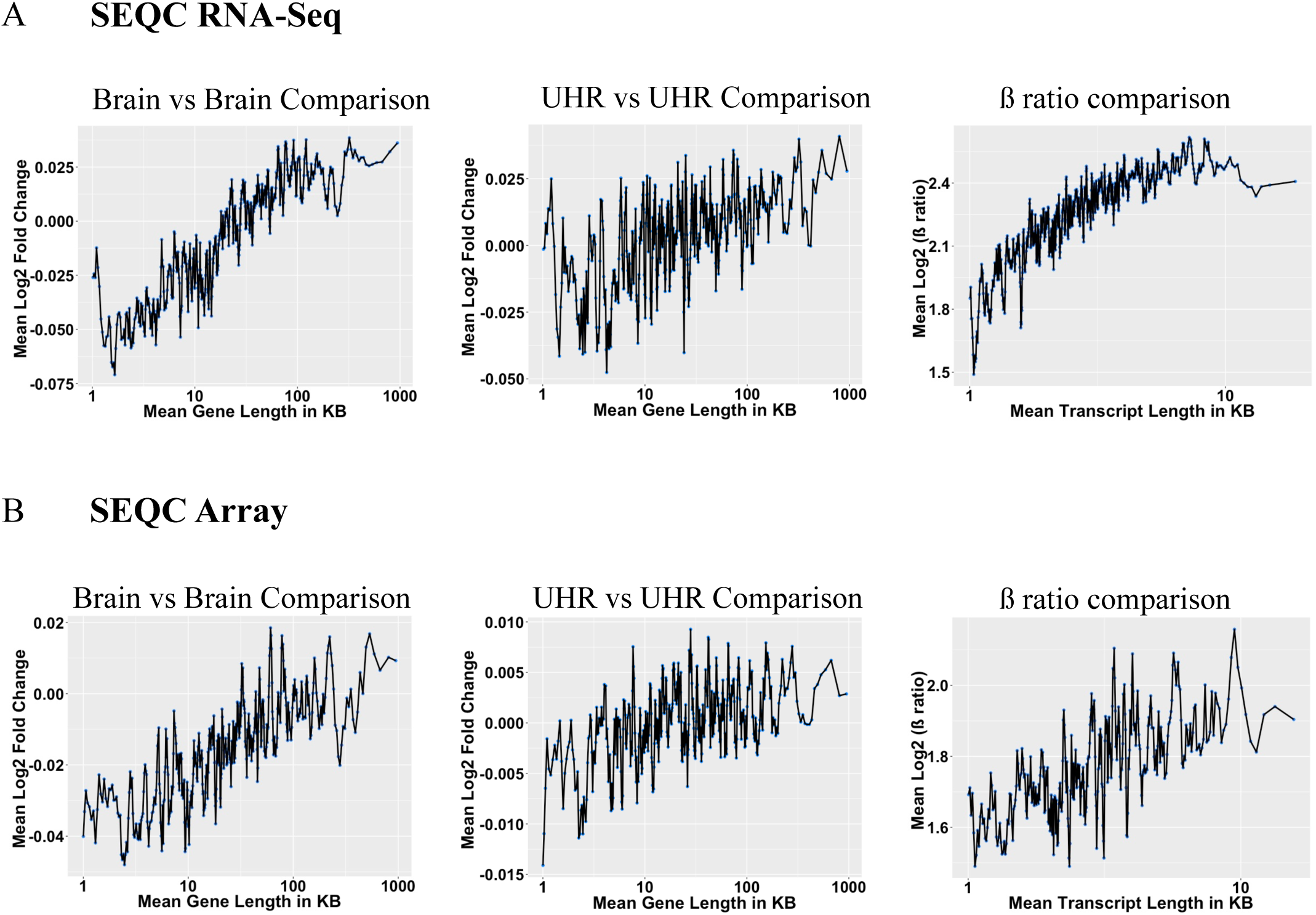
Long gene bias in the SEQC dataset (Related to Figure 4). **(A)** Brain vs. Brain randomized log fold-change plot against gene length (left panel; n = 32 each), Universal human reference (UHR) vs UHR randomized log fold-change plot against gene length (middle panel; n = 32 each) and log2 fold-change plot against transcript length using β ratio samples in RNA-Seq dataset (right panel; n = 32 each). **(B)** Brain vs. Brain randomized fold-change plot against gene length (left panel; n = 2 each), Universal human reference (UHR) vs UHR randomized log fold-change plot against gene length (middle panel; n=2 each) and log2 fold-change plot against transcript length using β ratio samples in microarray dataset (right panel; n = 2 each). Each blue dot is a bin of 200 genes with shift size of 40 genes (Gabel et al., 2015).

**Figure S7.**
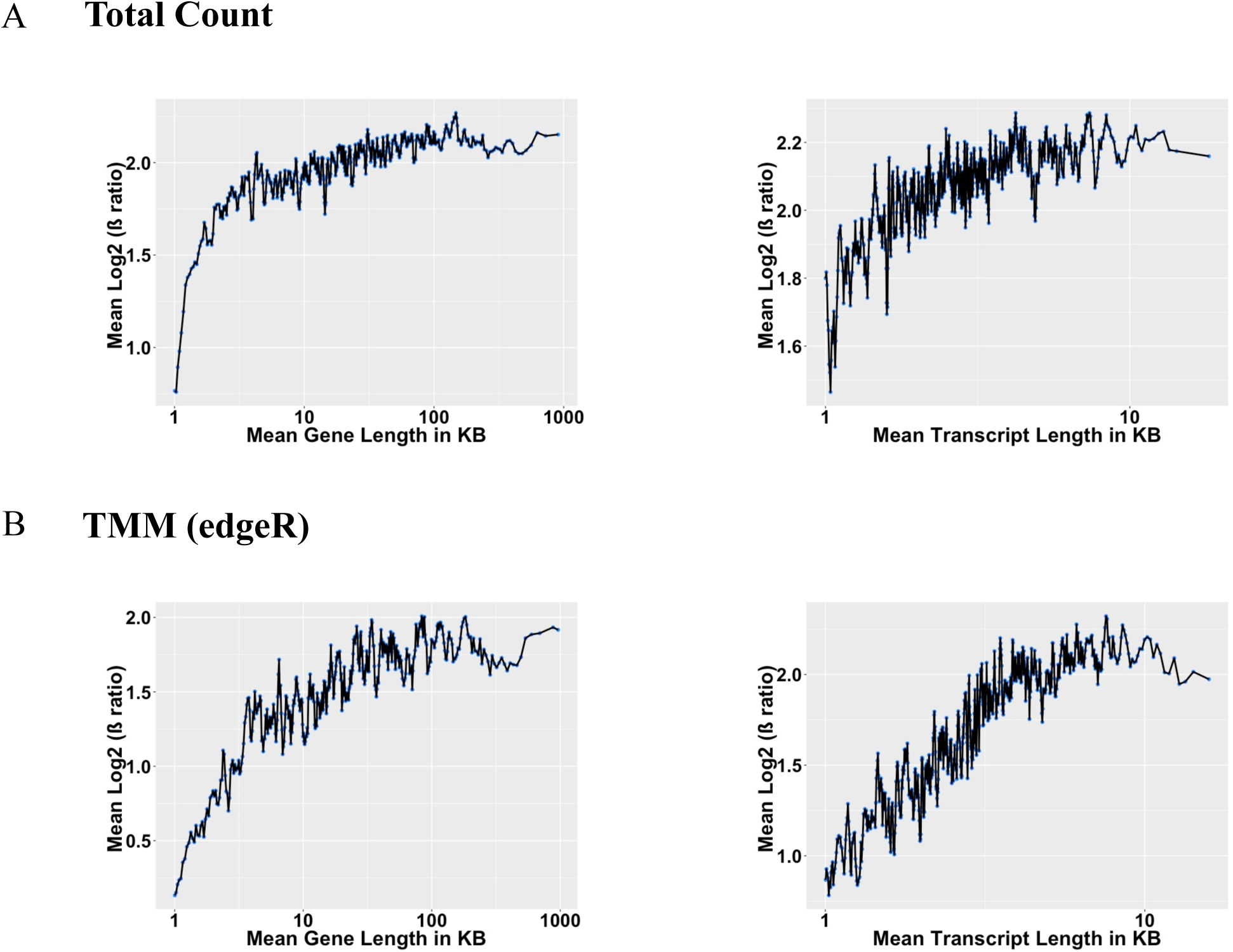
Long gene bias is independent of normalization methods (Related to Figure 4). Log2 fold-change plot against gene length using β ratio samples (n =64 each) for all genes using **(A) l**ibrary size normalization (or total count) against gene length (left panel) & transcript length (right panel) and **(B)** TMM (edgeR) normalization against gene length (left panel) & transcript length (right panel). Each blue dot is a bin of 200 genes with shift size of 40 genes (Gabel et al., 2015).

**Figure S8.**
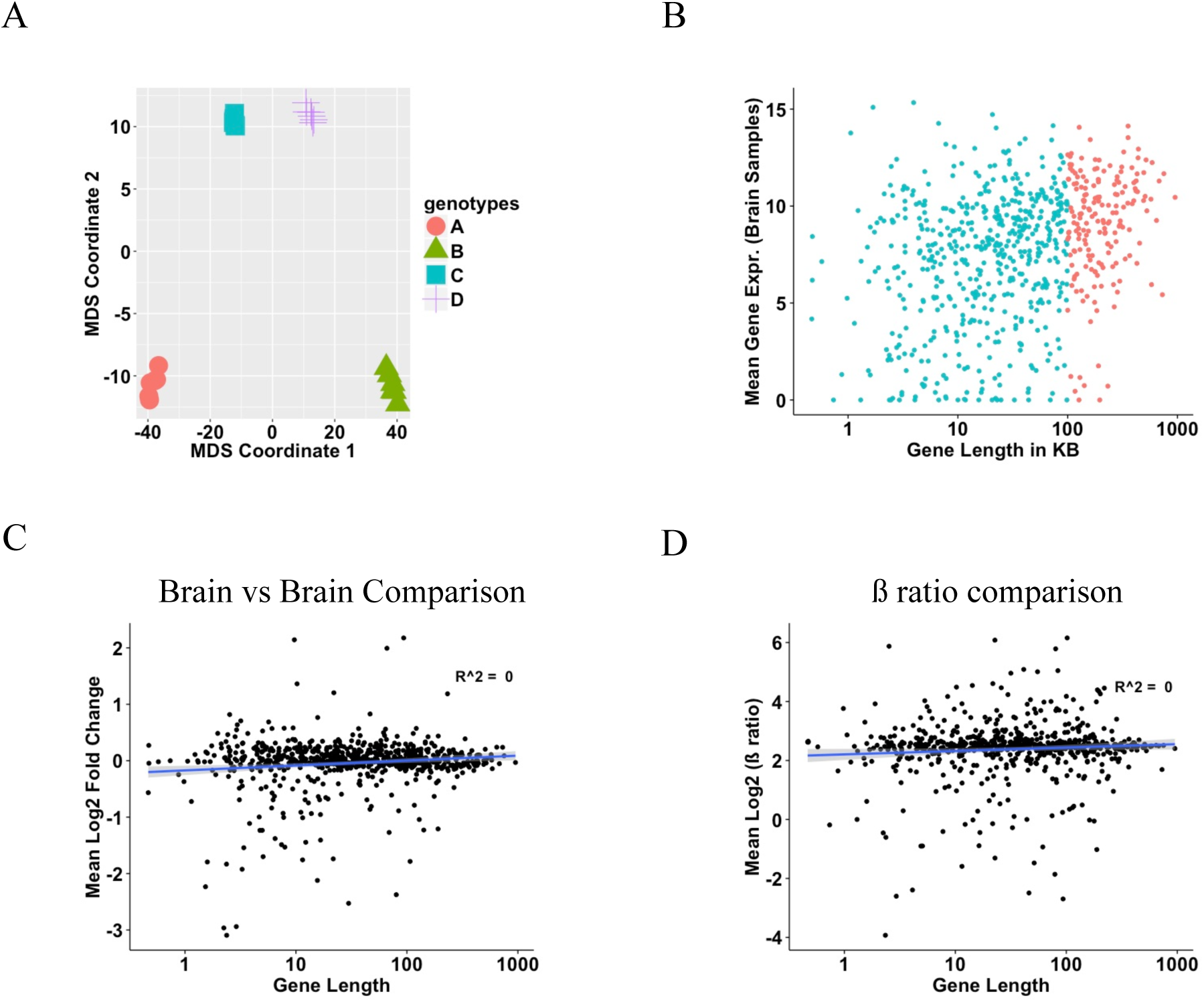
Long gene bias is not observed in Nanostring dataset (Related to Figure 4). **(A)** PCA plot on the NanoString dataset (n = 6 each sample type). **(B)** Scatter plot for mean gene expression in brain samples against its gene length **(C)** brain vs. brain randomized fold-change plot against gene length (n = 6 each). **(D)** Log2 fold-change plot against gene length using (B-A/C-A) = 4:1 samples (n =6 each).

**Figure S9.**
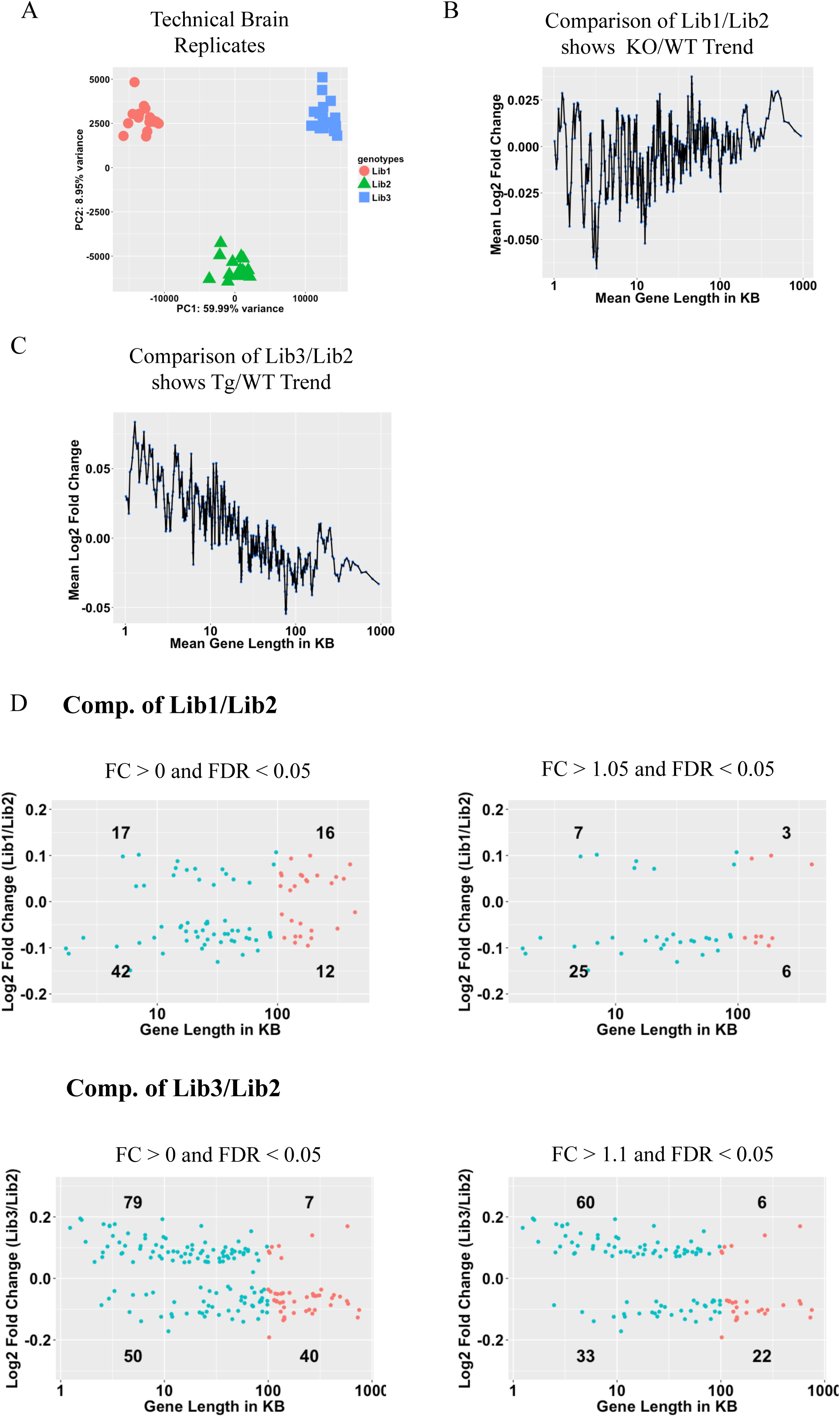
Explanation of reciprocal relationship among transcriptional changes between RTT and *MECP2* duplication syndrome. **(A)** PCA Plot of the B samples in Novartis SEQC dataset using library prep IDs. **(B)** Comparison of brain samples having library preparation 1 vs 2 against gene length (n = 16 each). **(C)** Comparison of brain samples having library preparation 3 vs 2 against gene length (n = 16 each). **(D)** Differential expression analysis between brain samples having library preparation id 1 vs 2 and 3 vs 2 across different fold changes and FDR < 0.05. Each blue dot is a bin of 200 genes with shift size of 40 genes (Gabel et al., 2015). The red and blue dot in (D) represent long and short genes, respectively.

**Figure S10.**
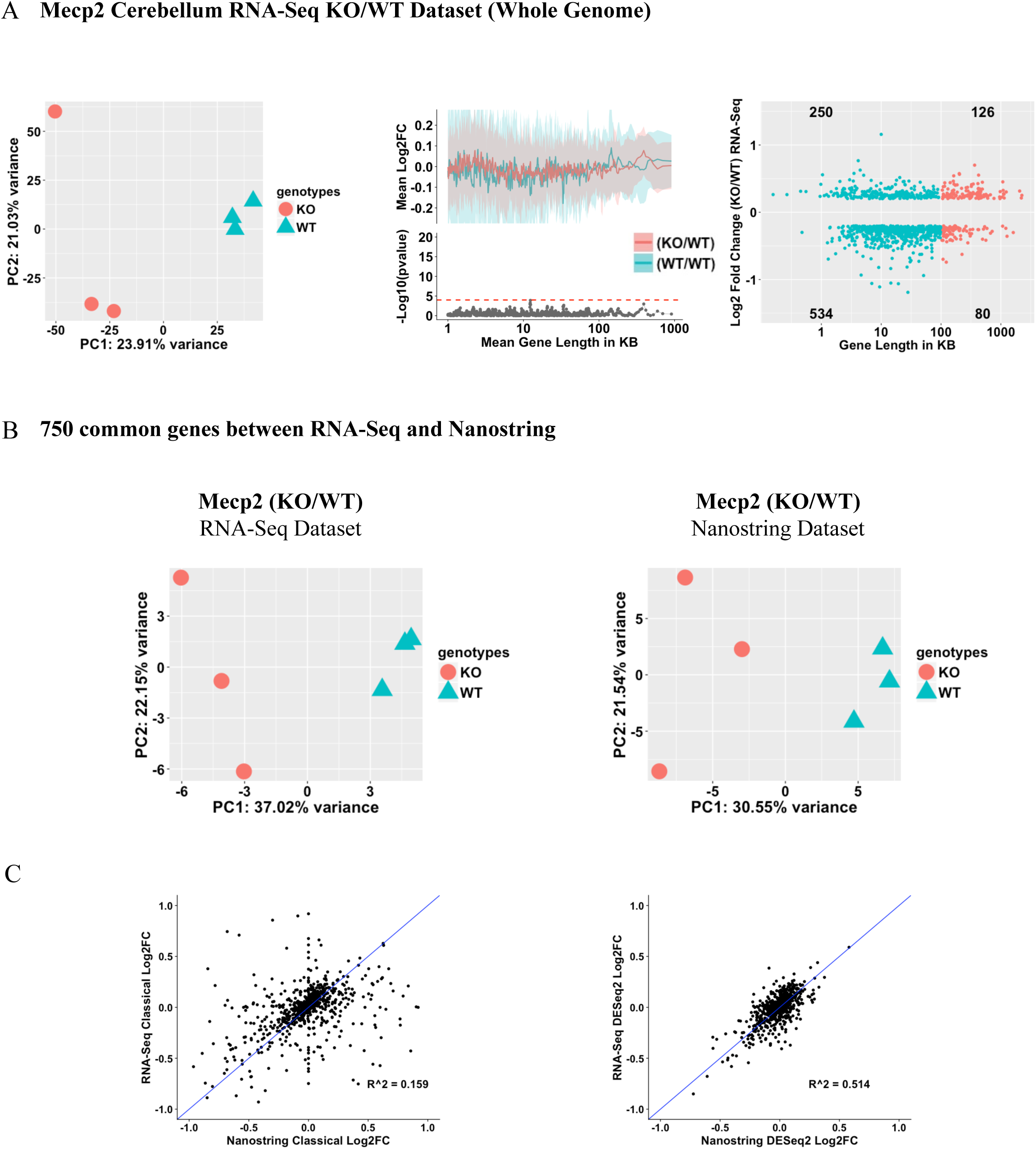
RNA-Seq and Nanostring analysis of Mecp2 KO and WT samples from the cerebellum of male mice (Related to Figure 5). A) Analysis using all the genes in the RNA-Seq cerebellum dataset. PCA Plot of Mecp2 KO and WT samples (left panel), overlap plot (middle panel) where, blue line (BL) represents the comparison of permuted WT/WT samples from a respective dataset (n = 1 each). The Red line (RL) represents the comparison of KO samples to its WT littermates (n = 3 each). The top half of each subgraph shows the lines that represent fold-change in expression for genes binned according to gene length (bin size of 200 genes with shift size of 40 genes) as described (Gabel et al., 2015). The blue and red ribbon correspond to one-half of one standard deviation of each bin for the comparison of WT/WT and KO/WT respectively. The bottom half of each subgraph is the p-value from the two-sample t-test between KO/WT and WT/WT. Bins with FDR < 0.05 are shown in red. The red dotted line indicates the minimum -Log_10_(p-value) that corresponds to a FDR < 0.05. Scatter plot of log fold-change in expression between KO and WT samples (right panel; n = 3 each) against gene length. The differentially expressed genes (FDR < 0.05 & absolute log2FC > log2(1.2)) were plotted. B) Analysis using 750 genes common in both RNA-Seq and Nanostring dataset. PCA plot of Mecp2 KO and WT samples (n = 3 each) by RNA-Seq (left panel) and Nanostring (right panel) platforms. C) Comparison of log2 fold changes using classical/standard method (left panel) and shrunken log2 fold changes (right panel) using DESeq2.

## STAR METHODS KEY RESOURCES TABLE

### Deposited Data

Table 1 has all details about the datasets used in the analysis with GEO Accession IDs. Software and Algorithms

Software and Algorithms
| Reagent or Resource | Source | Identifier |
| --- | --- | --- |
| STAR aligner (v2.4.2a) | Dobin et al., 2013 | https://github.com/alexdobin/STAR/releases/tag/STAR_2.4.2a |
| DESeq2 | Love et al., 2014 | https://bioconductor.org/packages/release/bioc/html/DESeq2.html |
| edgeR | Robinson et al., 2010 | https://bioconductor.org/packages/release/bioc/html/edgeR.html |
| NanoStringNorm | Waggott et al., 2012 | https://cran.r-project.org/web/packages/NanoStringNorm/index.html |
| GenomicFeatures | Lawrence et al., 2013 | https://bioconductor.org/packages/release/bioc/html/GenomicFeatures.html |
| ggplot2 | Hadley Wickham. ggplot2: Elegant Graphics for Data Analysis (2010) | https://github.com/tidyverse/ggplot2 |
| cowplot | CRAN Package | https://github.com/wilkelab/cowplot |

## CONTACT FOR REAGENT AND RESOURCE SHARING

Further information and requests for resources and reagents should be directed to and will be fulfilled by the Lead Contact, Zhandong Liu.

## EXPERIMENTAL MODELS AND SUBJECT DETAILS

### Mice

All mice used in this study were FVB.129 F1-hybrids. They were group-housed with up to five mice per cage. They were maintained on a 14h light:10h dark cycle (light on at 06:00) with standard mouse chow and water *ad libitum* in our AAALAS-accredited facility. All research and animal care procedures were approved by the Baylor College of Medicine Institutional Animal Care and Use Committee.

## METHOD DETAILS

### Analysis of Mecp2 datasets

The transcriptome datasets from *Mecp2* studies generated using microarray (GEO accession ids: GSE50225, GSE11150, GSE15574, GSE33457, GSE42895, GSE42987, GSE8720 and GSE6955) were downloaded from GEO. RMA function (Gautier et al., 2004; Irizarry et al., 2003) in the R “affy” package was used to perform background correction, normalization, and summarization of core probesets. NetAffx annotation files (Release 33 for mm9) was used to map affy probes to its official gene symbols. The expression values for genes with multiple probes were obtained by taking the average log2 expression value across all the probes corresponding to each gene. The NetAffx annotation file has information about the probe location, length and gene coordinates; we calculated gene length using the gene coordinates, and we specifically used gene length in all our figures where “Gene length in KB” is defined on the x-axis. We also ran our analysis on the transcript length (see figures S5A-B, right panel, and S6A-B, right panel). The extent of length-dependent bias with transcript length was similar to that of gene length. Since gene length information was not available in case of Affymetrix Human Genome U95 version 2 array, we mapped the probe to its gene and gene length using Ensembl Biomart database (version GRCh38.p5/Ensembl Genes 84).

The transcriptome dataset of the virtual cortex (Gabel et al., 2015) (GSE60077) was mapped to mm10 genome using STAR aligner v2.4.2a (Dobin et al., 2013) and for hypothalamus RNA-Seq dataset (Chen et al., 2015) (GSE66871), we used a published list of differentially expressed genes and normalized counts. For Johnson et al. (Johnson et al., 2017) RNA-Seq dataset (GSE83474), we used the raw count files provided by the authors in GEO. Similarly, for the transcriptome analysis of frontal and temporal cortex from RTT patients, we used the normalized gene expression profile provided in GSE75303 (Lin et al., 2016). We performed box plot and MDS plot to check for outliers in the sample distribution. The annotation files provided by GPL10558 were obtained to map Illumina probes to official gene symbols and RefSeq hg19 annotation was used to obtain gene length information.

### Running Average Plots

We used the same method as described in (Gabel et al., 2015) to compute the running average plot. In brief, the genes were sorted by their lengths and partitioned into bins using a sliding window of 200 consecutive genes in steps of 40 genes. The log_2_ fold-change values for genes within each bin were averaged. For consistency with the previous studies, we used genes whose lengths are between 1 kb and 1000 kb for all the plots. These plots were created using ggplot2 package in R.

### Confidence interval estimation in Overlap plots

We define the plots used in Figure 1 as “overlap plots,” meaning an overlap of two running average plots that shows intra-sample variation between control samples (WT) and inter-sample variation between two genotypes or conditions. To determine the amount of intra-sample variation, we computed the standard deviation of the genes in the same sliding window. By definition, 95% confidence interval for the mean is sample mean plus minus 1.96 times of the standard deviation. In all our overlap plots for the Mecp2 datasets, however, the confidence interval of KO/WT (or Tg/WT or D/V or RTT/WT) and WT/WT completely overlap. For the sake of legibility, we plotted only half of one standard deviation of the mean for each bin in the comparison of WT/WT and KO/WT (or Tg/WT or D/V or RTT/WT), which is denoted by the blue and red ribbon, respectively. Two-sample Student t-test was applied to each of the bins between KO/WT (or Tg/WT or D/V or RTT/WT) and WT/WT, followed by multiple hypothesis adjustment using the Benjamini-Hochberg method (FDR). The significant bins (FDR < 0.05) are denoted by red and non-significant bins are denoted by grey. The overlap plots were created using cowplot package in R.

### Distribution of differentially expressed genes in *Mecp2* datasets

To measure the distribution of long gene bias among differentially expressed genes, we extracted published lists of genes found to be significantly activated or repressed by *Mecp2* across different brain region. The published lists of differentially expressed genes were downloaded from the supplementary files in each study. Because of the frequent changes in gene name and annotation, we used MGI batch query (Eppig et al., 2015) to facilitate uniform comparison between these gene lists. The genomic locations were obtained for mm10/GRCm38. The original fold-change and FDR thresholds reported by respective publications were used. In case of microarray datasets, genes were plotted against their length. In the case of the RNA-Seq dataset, the calculation was done based on UCSC transcript IDs. Long genes (gene length > 100 Kb) were represented as red and short genes were represented as blue. The numbers of the upregulated and downregulated long/short genes are shown in four different quadrants.

### Analysis of SEQC dataset

We measured the long gene fold-change bias in RNA-Seq and microarray benchmark datasets, using the RNA-Seq datasets generated by all the Illumina HiSeq 2000 sites and microarray datasets generated by USF using Affymetrix Human Gene 2.0 ST Array in the SEQC consortium. The RNA-Seq raw count files and microarray PrimeView normalized file were accessed from the Gene Expression Omnibus database (GEO) (Barrett et al., 2013). The GEO accession IDs for the RNA-Seq and microarray datasets are GSE47774 and GSE56457, respectively. Raw count files from the Australian Genome Research Facility (AGR), Beijing Genomics Institute (BGI), Weill Cornell Medical College (CNL), City of Hope (COH), Mayo Clinic (MAY) and Novartis (NVS) were normalized using the DESeq2 method. Principal Component Analysis (PCA) and Multidimensional scaling plots (using Euclidean distance) were used to do a sanity check for a nominal amount of batch effects.

For further downstream analysis, we decided to use the Novartis dataset, as it had a minimal amount of non-biological variation (data not shown). The Novartis dataset consisted of 64 technical samples each of A (Universal Human Reference RNA), B (Human Brain Reference RNA), C (3A:1B) and D (1A:3B). We did not use sample type β (Ambion ERCC Spike-In Control Mix 1) or F (Ambion ERCC Spike-In Control Mix 2) in our analysis. For consistency with the SEQC consortium, we used hg19 iGenome NCBI/RefSeq annotation (build 27.2). The *transcripts* and *exon* functions in GenomicFeatures Bioconductor package (Lawrence et al., 2013) were used to obtain the gene and transcript length respectively, from the hg19 GTF file. Since a small number of genes or transcripts have multiple different genomic locations, genes or transcripts with the longest length were used. Expression values for genes with multiple transcript clusters were averaged across all transcript clusters corresponding to each gene. Similarly, for the microarray USF PrimeView dataset, sanity checks were performed using boxplot and MDS plots. Boxplots were used to check if the dataset was properly normalized and MDS plots were used to confirm that the dataset had a nominal amount of batch effects or non-biological variation.

### Library size normalization using Total Count and Trimmed Mean of M-values

To ensure that our normalization methods were not obscuring a genuine long gene bias, we normalized the raw counts from Novartis RNA-Seq dataset based on two other methods apart from DESeq2 (Love et al., 2014): a) Total Counts (Dillies et al., 2013) and b) the Trimmed Mean of M-values (TMM) method implemented in edgeR (Robinson et al., 2010; Robinson and Oshlack, 2010). For Total Counts, scaling factors were computed such that the normalized read counts across all samples are equal. In the case of the TMM method, we used the *calcNormFactors* function in the edgeR Bioconductor package to get the scaling factors and normalized read counts.

### SEQC NanoString sample preparation and analysis

We purchased Universal Human Reference RNA from Agilent Technologies, Inc., and Human Brain Reference RNA from Life Technologies, Inc. For the nCounter experiments, we used the same RNA sample types as SEQC. We assessed RNA purity and integrity with Bioanalyzer (Agilent Technologies, Inc.) prior to use in the nCounter assays. Sample preparation and analysis were done using a nCounter Prep Station 5s and a nCounter Digital Analyzer 5s. Expression of 770 genes (~730 genes with ~40 housekeeping genes and positive and negative controls) was assessed using the nCounter Human PanCancer Pathways Panel. A second PanCancer Pathways Panel was run using the same samples submitted to the first panel to assess the effect of batches on nCounter results. We used NanoStringNorm function (Waggott et al., 2012)in the R NanoStringNorm package to normalize the dataset. Boxplots and MDS plots were used for sanity checks. The two-sided Wilcoxon rank sum test was used to compare the distribution of the fold-change between long and short genes across the three different platforms.

### RNA isolation, sequencing and nanostring analysis from mouse cerebellum

We performed RNA extraction and purification from the cerebellum of male mice 8 to 9 weeks of age (three biological replicates of wild-type and Mecp2-null) using the Aurum™ Total RNA Fatty and Fibrous Tissue Kit (Bio-Rad 7326830) per the manufacturer’s instructions. Genomic DNA was eliminated using an on-column DNase digestion step. RNA quality was assessed using the Agilent 2100 Bioanalyzer system prior to library preparation for deep sequencing or use of the total RNA for Nanostring nCounter quantification.

RNA sequencing was performed using Illumina HiSeq 2000. All sequencing was done by the Genomic and RNA Profiling Core at the Baylor College of Medicine. For each sample, about 90 to 110 million pairs of 100 bp reads were generated. Raw reads were aligned to the *Mus musculus* genome (Gencode mm10; version M10) using STAR aligner v2.4.2a (Dobin et al., 2013) with default parameters. The overall mappability for all 6 samples was above 90% (Table 2). The read counts per gene were obtained using the *quantMode* function in STAR. These read counts are analogous to the expression level of the gene. Using the obtained raw counts, normalization and differential gene analysis were carried out using the DESeq2 package in the R environment. DESeq2 allows us to test for gene expression changes between samples in different conditions using more robust shrinkage estimation for dispersion and fold changes (Love et al., 2014). The default negative binomial generalized linear model with Wald test implemented in the package was used to identify significant differential expressed genes. Log fold change was calculated using both the classic method and shrinkage estimates calculated by DESeq2.

For the nCounter experiments, sample preparation and quality analysis were done using a nCounter Prep Station 5s and an nCounter Digital Analyzer 5s. Expression of 784 genes (750 endogenous genes with 34 housekeeping genes and positive and negative controls) was assessed using the nCounter Mouse PanCancer Pathways Panel. We used NanoStringNorm function (Waggott et al., 2012) in the R NanoStringNorm package to normalize the dataset and DESeq2 for differential expression analysis.

